# Diving efficiency at depth and pre-breeding foraging effort increase with haemoglobin levels in gentoo penguins

**DOI:** 10.1101/2023.05.14.539907

**Authors:** Sarah P. McComb-Turbitt, Glenn T. Crossin, Megan Tierney, Paul Brickle, Philip Trathan, Tony D. Williams, Marie Auger-Méthé

## Abstract

Individual differences in oxygen storage and carrying capacity have been associated with fitness-related traits and, for air-breathing aquatic animals, to diving ability and foraging success. In winter, many seabirds must replenish the energy reserves they have depleted during the breeding period. Thus, winter foraging efficiency can influence their upcoming breeding behaviour. Using gentoo penguins (*Pygoscelis papua*), we investigate (1) if inter-individual variation in diving efficiency (proportion of time spent at the bottom) is associated with indices of oxygen storage and carrying capacity (haemoglobin, haematocrit, body mass), and (2) if measures of pre-breeding foraging effort (mean trip duration, total time at sea, and vertical distance travelled) are associated with these oxygen indices and breeding status. Haemoglobin was positively correlated with diving efficiency, particularly for deeper dives, and only penguins with high haemoglobin levels frequently dove at depth ≥ 140 m. Such differences could affect resource access. However, potentially because reaching deep offshore waters requires travelling more than foraging nearshore, vertical distance travelled pre-breeding increased with haemoglobin levels. The relationship with haematocrit was non-linear, suggesting that commonly-used analyses may be inappropriate for this index. We found that early-laying penguins spent less time at sea prior to nesting than non-breeding penguins, suggesting that more efficient foragers lay earlier. Given that diving efficiency at depth is linked to aerobic capacity, anthropogenic activities taking place in either nearshore or offshore waters (e.g., shallow water fisheries, offshore oil rigs) may have differing impacts on individuals. Further understanding these links could help the conservation of diving species.

## 1. Introduction

Intra-specific variation in foraging and movement behaviour is ubiquitous, and can have important ecological consequences at both the individual and population levels (Bolnick et al. 2003, Ceia & Ramos 2015, Phillips et al. 2017, Shaw 2020). Such variation can be caused by complex relationships with the environment, intra- and inter-specific interactions (e.g., competition), and physiological differences (Shaw 2020). Differences in oxygen storage and carrying capacity can be particularly important, as they have been associated with variation in movement behaviour, habitat selection, and fitness-related traits such as annual breeding success and arrival at breeding grounds (Crossin et al. 2013, 2015, Minias 2015). For diving animals, oxygen storage and carrying capacity can limit diving ability, and thus access to resources may be physiologically restricted (Roncon et al. 2018). For such animals, the time spent diving and the rate of mass gain have been linked to indices of aerobic capacity (Crossin et al. 2015).

Diving animals have a range of adaptations to maximize their aerobic capacity and extend their breath-holding capability (Roncon et al. 2018). Haemoglobin is an oxygen-carrying protein present in red blood cells, and thus its level is a crucial determinant of the rate of oxygen delivered to tissues (Minias 2015). Haemoglobin levels are often used as a proxy of aerobic capacity and physiological conditions in birds (Minias 2015), and is elevated in bird species that dive (Minias 2020). Prolonged breath-holding capacity is often linked to oxygen storage. Oxygen stores typically increase with haemoglobin concentration, blood volume, myoglobin concentration in skeletal muscles, muscle mass, and lung volume (Mirceta et al. 2013, Roncon et al. 2018). As larger individuals possess more blood and muscle, body mass is often positively associated with oxygen storage capacity (Ponganis & Kooyman 2000, Cook et al. 2013, Mirceta et al. 2013, Polito et al. 2015, Camprasse et al. 2017). In addition, consistent with generally observed allometric patterns of metabolism, larger body mass is often associated with lower oxygen usage per unit mass (Glazier 2005, Hudson et al. 2013). As such, total blood haemoglobin concentration (referred as Hb), haematocrit (percentage of red blood cells in blood, referred as Hct), and body mass can be important determinants of a breath-holding diver’s physiological performance and can limit an animal’s diving ability (Crossin et al. 2015, Chimienti et al. 2017).

Seabirds display high levels of individual variation in foraging and movement (Ceia & Ramos 2015, Phillips et al. 2017), and diving seabirds provide a unique opportunity to link differences in oxygen storage and carrying capacity to variation in diving and movement behaviour, and ultimately to differential breeding participation. Many seabirds severely deplete their energy reserves during breeding and moulting periods and must replenish these reserves by foraging intensively in winter (Sorensen et al. 2009, Crossin & Williams 2016). Consequently, how efficient an individual is in acquiring food in winter can influence multiple aspects of its breeding capacity and behaviour (Daunt et al. 2006, Shoji et al. 2015). For example, individuals that spend less time foraging in winter, and thus are assumed to be more efficient foragers, breed earlier (Daunt et al. 2006). Understanding how individuals vary in their foraging efficiency and behaviour as they prepare for breeding can help assess the possible impacts natural and anthropogenic ecosystem changes can have on populations (Durell 2000, Bearhop et al. 2006, Cury et al. 2011, Phillips et al. 2017).

Here, using a population of gentoo penguins (*Pygoscelis papua*), we explore how individual variation in winter diving behaviour relates to indices of oxygen storage and carrying capacity, and how these relate to their breeding status. We use gentoo penguins because they show considerable individual variation in winter movement behaviour (Baylis et al. 2021). Gentoo penguins consume a wide variety of locally available prey found at a range of depths down to approximately 200 m (Tanton et al. 2004, Thiebot et al. 2011, Camprasse et al. 2017). While they are considered a generalist species, many individuals have a specialized diet and foraging behaviour (Polito et al. 2015, Camprasse et al. 2017, Handley et al. 2017, Herman et al. 2017). There is a high likelihood of intraspecific competition for prey around colonies, as gentoo penguins are not migratory and thus forage year-round relatively close to their breeding colonies (Clausen et al. 2005, Bearhop et al. 2006, Kokubun et al. 2010). In addition, given their magnitude, certain commercial fisheries in the Falkland Islands could reduce the availability of important prey species in winter (Clausen & Pütz 2003). The number of breeding pairs in the Falkland Islands fluctuates drastically, with changes of 50 % occurring in a single year (Pistorius et al. 2010, Stanworth & Crofts 2019). Changes in population breeding success are thought to be linked to prey availability (Pistorius et al. 2010) and broad-scale climatic variation (Baylis et al. 2012).

To understand the link between oxygen storage and carrying capacity, pre-breeding diving and foraging behaviour, and breeding activity, we investigate (1) if diving efficiency (proportion of the time within a dive spent at the bottom) is associated with indices of oxygen storage and carrying capacity (Hb, Hct, body mass), and (2) if measures of pre-breeding foraging effort (mean trip duration, total time at sea, and vertical distance travelled) differ between individuals based on their breeding status and indices of oxygen storage and carrying capacity. We use time-depth recorders (TDRs) to quantify diving efficiency and foraging effort. As attaching such devices to animals can add drag and affect their performance, behaviour, and reproductive success (Taylor et al. 2001, Beaulieu et al. 2010, van der Hoop et al. 2014, Wilson et al. 2015), we also evaluate whether tagged individuals differ from control birds in terms of breeding activity, Hb, Hct, and mass.

## 2. Methods

### 2.1 Data collection

We captured 66 adult gentoo penguins post-moult (April 4 – 10, 2018) from two colonies in the Falkland Islands: Rookery Sands, Race Point (51.4345°S, 59.1081°W, n=33) and Tamar Point, Pebble Island (51.3241°S, 59.4523°W, n=33). We weighed each captured penguin using a 10 kg Pesola scale, took a blood sample (< 1.5 ml) from the left brachial vein using heparinized syringes fitted with 25-gauge needles, and marked them with black hair dye (L’Oreal Preference) to facilitate re-identification. We equipped each penguin with a Sirtrack K2G 173A SWS KIWISAT 202B Argos tag (34g, 55 x 27 x17 mm, Sirtrack, Havelock North, New Zealand) and a Lotek LAT1800 TDR tag (13.6 g, 62 x 13 x 13 mm, Lotek Wireless Inc, St. John’s, Canada). As in Handley et al. (2018) and Baylis et al. (2021), we attached the two devices to midline back feathers using overlapping layers of TESA^®^ tape (Beiersdorf AG, GmbH, Hamburg, Germany) and sealed the tape seams with cyanoacrylate glue (Supplement 1).

During the breeding season (September 27 - October 21, 2018), we recaptured 35 of the previously tagged penguins and 35 unmarked penguins, the latter serving as controls in our analysis of the potential effects of tagging (Table 1). As described above, we weighed each penguin, took a blood sample, and marked the control bird with hair dye. We retrieved any devices still attached to the penguins. Twenty-one penguins had TDR devices still attached (7 Race Point, 14 Pebble Island, Table 1), one of which still had an Argos tag attached (from Race Point). We note that we were unable to collect sufficient blood samples from two individuals with a TDR tag, resulting in only 19 of the individuals with TDR data also having blood data from the breeding period (5 Race Point, 14 Pebble Island, Table 1).

**Table 1.**
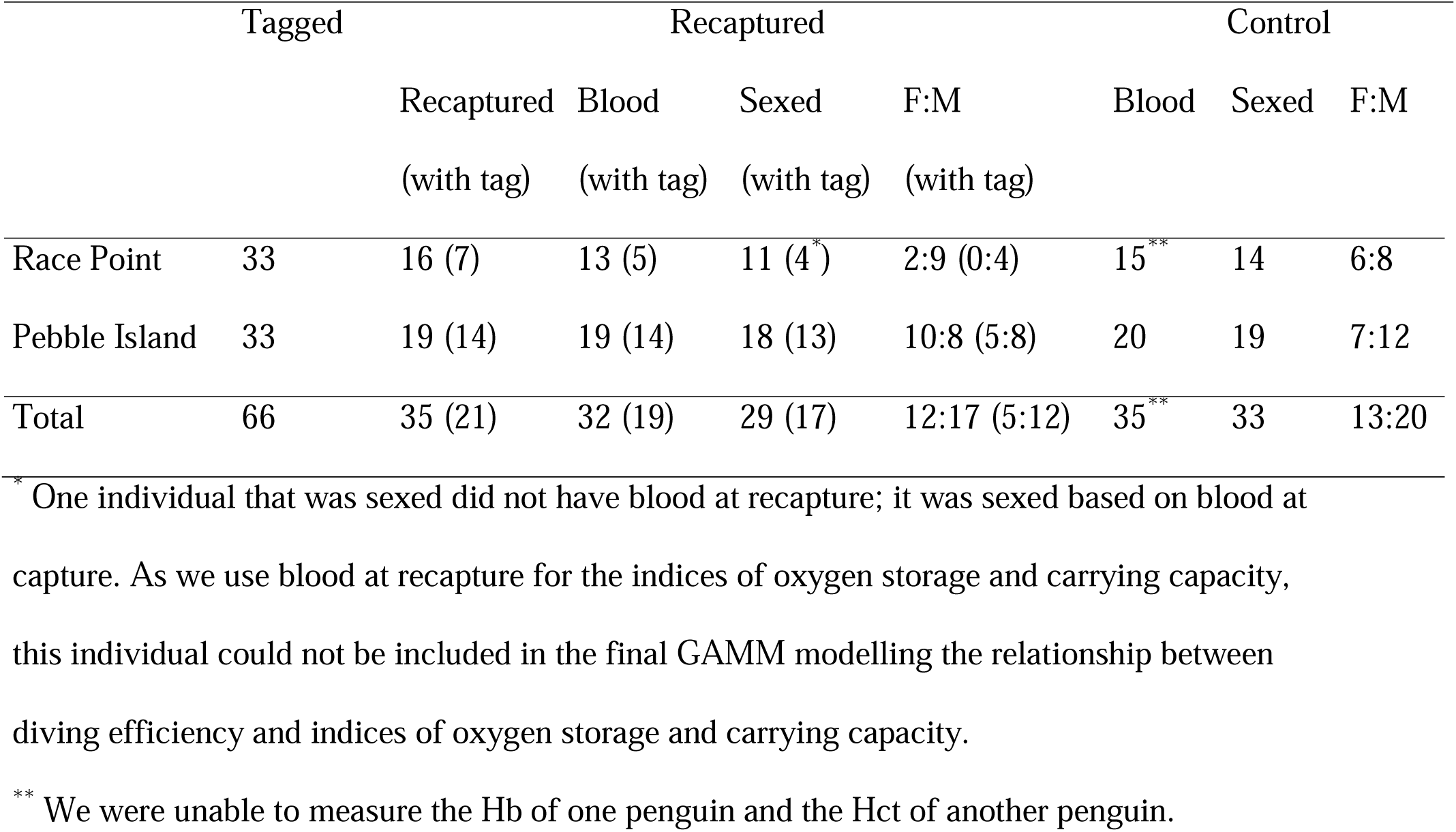
The number of individuals tagged, recaptured, and sampled as control. We state how many individuals were captured and recaptured at each colony, the number of individuals for which we had blood samples, the number that could be sexed, and the female-to-male sex ratio (F:M). For the recaptured penguins, we present in brackets the number of individuals for which the TDR tags were still attached at recapture.

|  | Tagged | Recaptured |  |  |  | Control |  |  |
| --- | --- | --- | --- | --- | --- | --- | --- | --- |
|  |  | Recaptured<br>(with tag) | Blood<br>(with tag) | Sexed<br>(with tag) | F:M<br>(with tag) | Blood | Sexed | F:M |
| Race Point | 33 | 16 (7) | 13 (5) | 11 (4<sup>*</sup>) | 2:9 (0:4) | 15<sup>**</sup> | 14 | 6:8 |
| Pebble Island | 33 | 19 (14) | 19 (14) | 18 (13) | 10:8 (5:8) | 20 | 19 | 7:12 |
| Total | 66 | 35 (21) | 32 (19) | 29 (17) | 12:17 (5:12) | 35<sup>**</sup> | 33 | 13:20 |

<sup>\*</sup> One individual that was sexed did not have blood at recapture; it was sexed based on blood at capture. As we use blood at recapture for the indices of oxygen storage and carrying capacity, this individual could not be included in the final GAMM modelling the relationship between diving efficiency and indices of oxygen storage and carrying capacity.
<sup>\*\*</sup> We were unable to measure the Hb of one penguin and the Hct of another penguin.

The research was conducted under Falkland Islands Scientific Research Licence (R12/2017) and conformed to guidelines from the Canadian Committee on Animal Care (University of British Columbia Animal Care Permit A17-0243 and Dalhousie University Animal Care Permit 17-100).

### 2.2 Indices of oxygen storage and carrying capacity

Using the blood samples from the breeding period, we used haematocrit (Hct) and haemoglobin (Hb) as physiological indicators of blood oxygen storage and carrying capacity. As Hct is based on blood cytology and Hb is based on blood biochemistry, each measure provides different insights into the aerobic capacity of individuals (Fair et al. 2007, Kaliński et al. 2011, Minias 2015). Hct (percent packed red blood cell volume) was determined from fresh whole blood in heparinized capillary tubes centrifuged for 5 min (Centrifuge; Brinkmann Instruments, Ontario, Canada) at 10000 g and measured using digital callipers (± 0.01 mm). Hb (total blood haemoglobin concentration in g/dL) was determined using the cyanmethemoglobin method (Drabkin & Austin 1932), where 5 μl of fresh whole blood was pipetted (Gilson Pipetman P2; Gilson Medical Electronics, France) into 1.25 ml Drabkin’s reagent (D541-6V, Brij 35 solution B4184-100ml; Sigma-Aldrich, Missouri, US). We ran triplicate samples and read absorbance at 540 nm with a microplate spectrophotometer (BioTek PowerWave 340, Vermont, USA). The inter-assay coefficient of variation was 1.70 % between plates.

We used the body mass measured during the breeding period as a proxy of oxygen storage capacity. We did not scale body mass to structural body size to generate a morphometric index because (1) oxygen stores (e.g., blood volume) and dive duration have been shown to increase with body mass itself (e.g., Hassrick et al. 2010), and (2) mass alone has been shown to be a more reliable measure of body condition in birds (Schamber et al. 2009, Labocha & Hayes 2012). We identified the sex of the individuals by sending blood samples to Viaguard^†^, which performs DNA testing. We note that sex results were unavailable for three of the 19 penguins for which we had TDR and blood data (Table 1).

### 2.3 Dive data processing

The TDR tags recorded depth (0.25 m resolution), temperature (0.01°C resolution), and wet/dry state at 1 s intervals. We wanted to focus on the time period that could be linked to the blood samples. Since the maximum life span of red blood cells in birds is estimated to be 35 – 45 days (Rodnan et al. 1957), we limited the analyses to dives made within 40 days prior to recapture. Given that the last Argos transmission was August 16, 2018, well before this threshold, and very few tags transmitted locations past May, we do not consider the Argos data further.

To characterise the diving behaviour of foraging individuals, we extracted the dive metrics from the TDR series using the software Divebomb (1.0.7, Nunes 2019) in Python (3.4.10, Python Software Foundation 2019). We first corrected the dive data for drift using a zero offset (Luque & Fried 2011), and we identified the following dive metrics: start of the dive, dive duration, maximum depth, bottom time, and post-dive surface interval. To remove the shallow dives most likely associated with travelling, we discarded all dives shallower than 5 m (Kokubun et al. 2010, Lee et al. 2015, Carpenter-Kling et al. 2017). Following Kokubun et al. (2010) and Lee et al. (2015), we eliminated the inclusion of long surface periods unlikely to be associated with active foraging by only considering dives with post-dive surface interval < 200 s. Through a visual assessment of 100 random dives across all individuals, we quantified that Divebomb gave precise measurements of decent, bottom, accent, and surface phases of dives 94 % of the time. Divebomb produced minor errors distinguishing the bottom phase in cases where undulations in depth occurred over the transition from decent to accent phases.

### 2.4 Behavioural observations

To assess the reproductive status of penguins, teams of two observers monitored the breeding behaviour of marked birds. Through daily colony scans and the continuous monitoring of arriving individuals, we identify which individuals were present at Race Point from September 27, 2018, to October 24, 2018, and Pebble Island from September 29, 2018, to October 24, 2018. Between 13h00 – 19h00 daily, we conducted focal follows on each individual observed in the colony. The focal follows were at least 5 min long and lasted until the presence or absence of eggs in an occupied nest could be confirmed. The pair bonding behaviours we monitored included nest building, copulation, pair calling, and pair bowing (Williams 1995). Gentoo penguins in the Falkland Islands lay their first egg from mid-October to early November (Otley et al. 2004, Black et al. 2018). As we left the field site before we could acquire the lay date of many penguins, we classified penguins into three categories of reproductive status: non-breeding, breeding, and early-laying individuals. Penguins classified as non-breeders were found most often at different locations than nesting penguins, typically towards the edge of colonies, and rarely displayed pair-bonding behaviours. Penguins categorized as breeders were observed conducting pair-bonding behaviours at stable nest site locations. Early-laying penguins were observed to be incubating at least one egg prior to October 24, 2018. We note that we had individuals of both sexes in each reproductive category and that none of the penguins had laid an egg at Race point at the time of departure of field observers.

### 2.5 Linking diving efficiency to oxygen storage and carrying capacity indices

We used a generalized additive mixed model (GAMM) to determine the relationship between diving efficiency at different maximum depths and our three previously described indices of oxygen storage and carrying capacity (Hb, Hct and body mass). We defined diving efficiency as 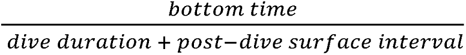 (Simeone & Wilson 2003, Lescroël & Bost 2005, Lee et al. 2015). Using the R package mgcv (Wood 2011), we used a thin-plate regression spline to model the non-linear relationship between diving efficiency and maximum dive depth. To investigate the potential interactions between maximum depth and the indices of oxygen storage and carrying capacity (Hb, Hct, and mass), we included a tensor product term (with cubic regression spline basis) between maximum depth and each index. To be able to decompose the interactions with maximum depth from the potential main effects of each index, we included a separate tensor product and a thin-plate regression spline for the relationship with diving efficiency and each index (i.e., used s() + ti() in mgcv). To account for potential biological or behavioural differences in sex and colony, we first included both terms as fixed effects in our GAMM and assessed their importance. As sex was not deemed an important covariate for this analysis (see Supplement 2) and three individuals were missing sex information, our final analysis did not include sex and included these three individuals. Thus, our final GAMM was applied to the data of 19 individuals. We also included an individual ID random effect on the intercept to account for the fact that we had multiple dives per penguin.

The diving data were highly temporally autocorrelated, and fitting GAMMs with complex autoregressive-moving-average (ARMA) functions are computationally demanding. For example, running our full GAMM with an ARMA function took multiple days to run on an iMac 3.6 GHz 10-Core Intel Core i9, and we ran into multiple convergence problems. Thus, we assessed how much sub-sampling was required to remove the autocorrelation in the residuals of the full model and found that we had to keep only one dive every 70 dives to completely remove the autocorrelation in the residuals. We used this 70-dive sub-sampling scheme, which resulted in having on average 85 dives per individual. To identify which of the covariates of the GAMM should be selected, we used the shrinkage method with a double penalty (i.e., select = “TRUE” in mgcv; Marra & Wood 2011), which penalizes both unnecessary wiggliness and linear effect of a covariate. This approach has many advantages over stepwise variable selection and model subset selection, such as not limiting the combinations of covariates explored, and has been previously used to select covariates in diving models (Marra & Wood 2011, Photopoulou et al. 2014).

There was some correlation and concurvity between the covariates. As such, we explored whether, and how, the results changed when creating GAMMs with only one of the important oxygen indices at a time. See Supplement 3 for details. In addition, the residuals of the final GAMM displayed evidence of heteroskedasticity. This variation in variance can be attributed to the fact that our final GAMM modelled proportion data (as defined, diving efficiency ranges between 0 and 1) with a normal error distribution. As such, we also explored whether, and how, the results changed if we used a GAMM with a beta regression. Supplement 4 details how the results remained largely unchanged.

As post-hoc analyses, we further explored how the deep diving behaviour of individuals changed in relation to the oxygen storage and carrying capacity indices deemed important in the GAMM above. As some individuals with low oxygen storage and carrying capacity may reduce, or altogether forgo, doing deep dives (here defined as ≥ 140 m), we used simple generalized additive models (GAMs) to assess whether the proportion of deep dives each individual performed changed in relation to each of the selected oxygen indices. As high diving efficiency can be achieved by either extending the bottom time or reducing surface recovery between dives, we used new GAMMs to explore how bottom time and post-dive surface intervals for deep dives changed in relation to each of the selected oxygen indices. For these new GAMMS, we looked at the oxygen indices separately, as the smaller sample size did not allow to have both in the models at once. We included an individual ID as a random effect to account for the fact that we have more than one dive per penguin.

### 2.6 Associating measures of foraging effort with breeding status and oxygen storage and carrying capacity indices

To determine if pre-breeding foraging behaviour changes between individuals based on their breeding status and oxygen storage and carrying capacity, we investigated three measures of foraging effort: mean trip duration, time at sea, and vertical distance travelled. To estimate mean trip duration and time at sea, we first identified trips at sea using the binary wet-dry data collected with the saltwater sensor of the TDR tags. To account for occasional jumps at sea, which results in brief recordings of dry conditions, we averaged the wet-dry data using a 10-minute rolling window. We classified any window value with an average > 0 as wet. We terminated a trip just before a dry period lasting at least one hour, and the subsequent trip began once a new wet value was recorded. The mean trip duration was the average duration of trips for the 40 days prior to recapture, while the time spent at sea was the sum of all trip durations within that period. Our last measure of foraging effort was the total amount of vertical distance travelled for the time spent at sea (*∑ maximum depth (km)* × 2; similar to Lescroël & Bost 2005, Booth et al. 2018).

Using multiple linear regressions, we investigated whether each of these measures of foraging effort (mean trip duration, time at sea, and vertical distance travelled) was associated with breeding status, sex, and the three oxygen storage and carrying capacity indices (Hb, Hct, mass). We only included individuals from Pebble Island since none of the Race Point individuals laid eggs before the end of the field season. To identify the covariates that best explained the data, we fitted models with all possible combinations of three of these covariates. Given the small sample size (n=13), we compared the models using the Akaike Information Criterion with correction for small sample size (AICc). We considered models that outperformed the null model (model with no covariates) and had ΔAICc < 4 from the best model (Burnham and Anderson 2002). Next, we used Wilcoxon rank sum tests to assess whether there were differences in oxygen indices across sex and breeding status. We used a Bonferroni correction to account for the multiple comparisons.

### 2.7 Assessing the potential effects of tagging

To limit the potential influence of our tags on the behaviour, energy expenditure, or condition of the animals (e.g., Wilson et al. 1986, Vandenabeele et al. 2015), we minimized drag by using tags with a streamlined design and attaching them in an appropriate caudal position (Bannasch et al. 1994). Furthermore, the overall weight of devices was < 1 % of the average body mass recorded, and similar devices have not affected the diving behaviour of gentoo penguins (Kokubun et al. 2010). However, especially because we deployed tags for an extended period, we wanted to assess whether the data for the tagged penguin were representative of the untagged animals and look for signs of tagging effects. To do so, we compared the physiology at breeding and breeding status of tagged to untagged penguins. Specifically, we used Welch two-sample t-tests to compare Hb, Hct, and mass at breeding. For the reproductive status, we used χ^2^ goodness-of-fit tests, and because of the small sample size (37 individuals for 6 categories), we estimated the p-value using a Monte Carlo method (Hope 1968). We note that we were unable to sample sufficient blood for three of the recaptured penguins, as well as unable to measure Hct for one untagged penguin and Hb for another untagged individual. Thus, while the mass comparison included 35 tagged, and 35 untagged individuals, the comparisons for Hct and Hb included only 32 tagged and 34 untagged individuals. In addition, for the comparison of reproductive status, we focussed only on penguins from Pebble Island since none of the penguins at Race Point laid an egg before the end of our field season. Finally, we also quantify the proportion of tagged penguins that increased in mass between capture and recapture and the average weight gain during that period.

All data analyses were performed using R (4.2.1, R Core Team 2022). Values presented in the results are mean ± SD unless otherwise stated, and statistical tests were considered significant at a 0.05 level. Supplement 8 provides the code and data needed to reproduce the analyses.

## 3. Results

### 3.1 Relationships between diving efficiency and indices of oxygen storage and carrying capacity

Our GAMM indicated that diving efficiency had statistically significant relationships with colony, as well as with the maximum depth of a dive, and its interaction with Hb and Hct (Table 2, Fig. 1). Diving efficiency was highest at intermediate depths followed by a decline at deeper depth (Fig. 1A). The interaction between maximum depth and Hb showed that while penguins with higher Hb did not appear to have higher diving efficiency for shallow dives, diving efficiency increased with Hb for deeper dives (Fig. 1B). In the post-hoc analyses, the relationship between bottom time for deep dives and Hb was positive and statistically significant, while the relationship between Hb and post-dive surface interval of deep dives was not statistically significant (Table 3). The results of these post-hoc analyses indicate that this increase in diving efficiency with Hb at deeper depth is primarily explained by an increase in bottom time rather than a decrease in the post-dive surface interval (Fig. 2B-C). In addition, we found that the proportion of deep dives (≥ 140 m) made by an individual increased with Hb (Fig. 2A), a relationship that was statistically significant (Table 3).

**Figure 1.**
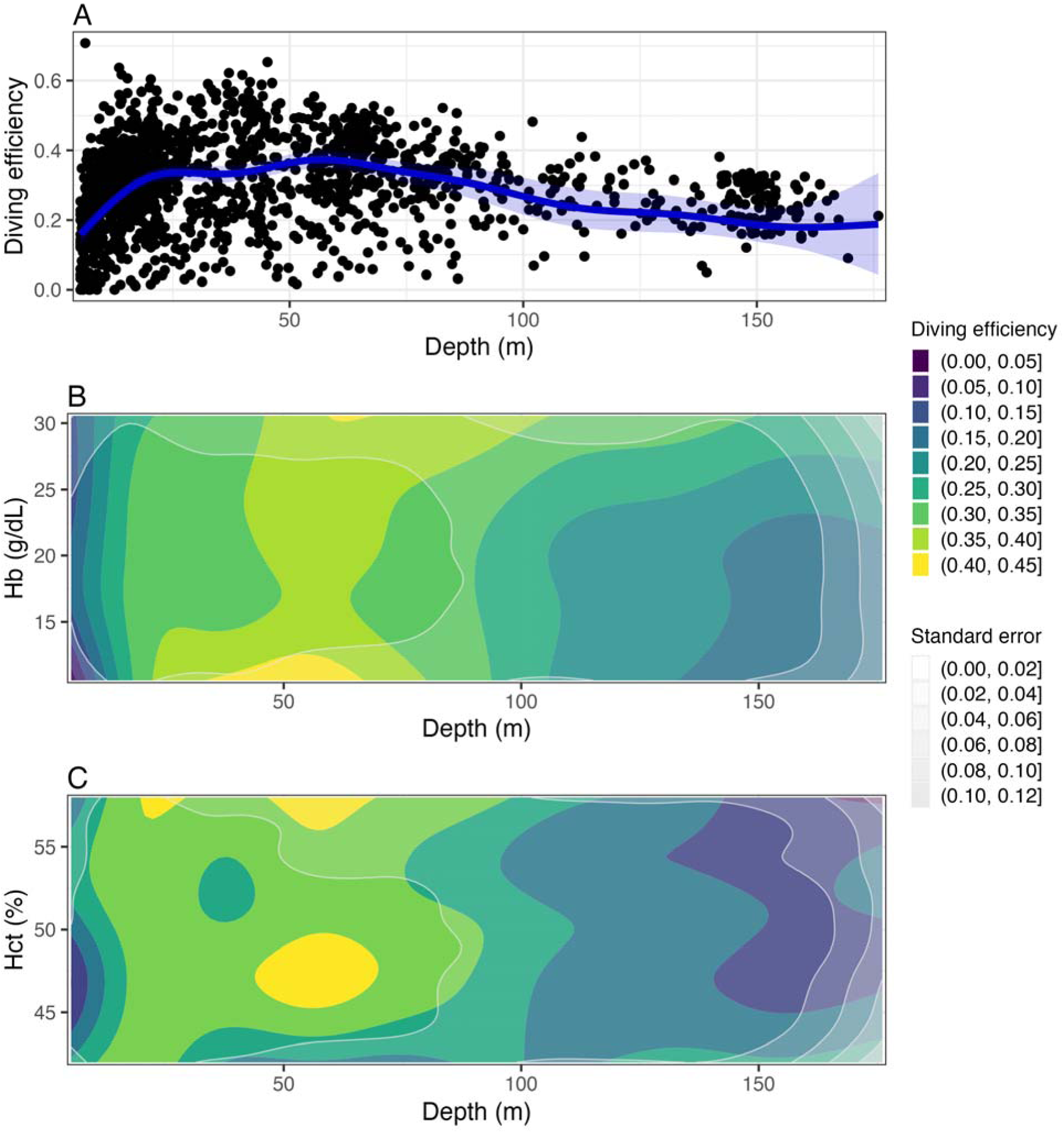
Slices of predictions from the final GAMM. Panel A displays how the predicted diving efficiency changes in response to varying maximum depth values of a dive. The blue band represents the 95% confidence intervals. The points show the observed values. Panel B shows how diving efficiency is predicted to change with both depth and Hb, while panel C shows how diving efficiency is predicted to change with both depth and Hct. For both these panels, the colour represents the diving efficiency value (yellow representing highest efficiency) and the grey bands represent the standard error (more opaque grey representing higher standard error). The areas with larger standard error values (i.e., more opaquely shaded) should be interpreted more cautiously. A total of 19 penguins were included in this analysis (5 from Race Point and 14 from Pebble Island).

**Figure 2.**
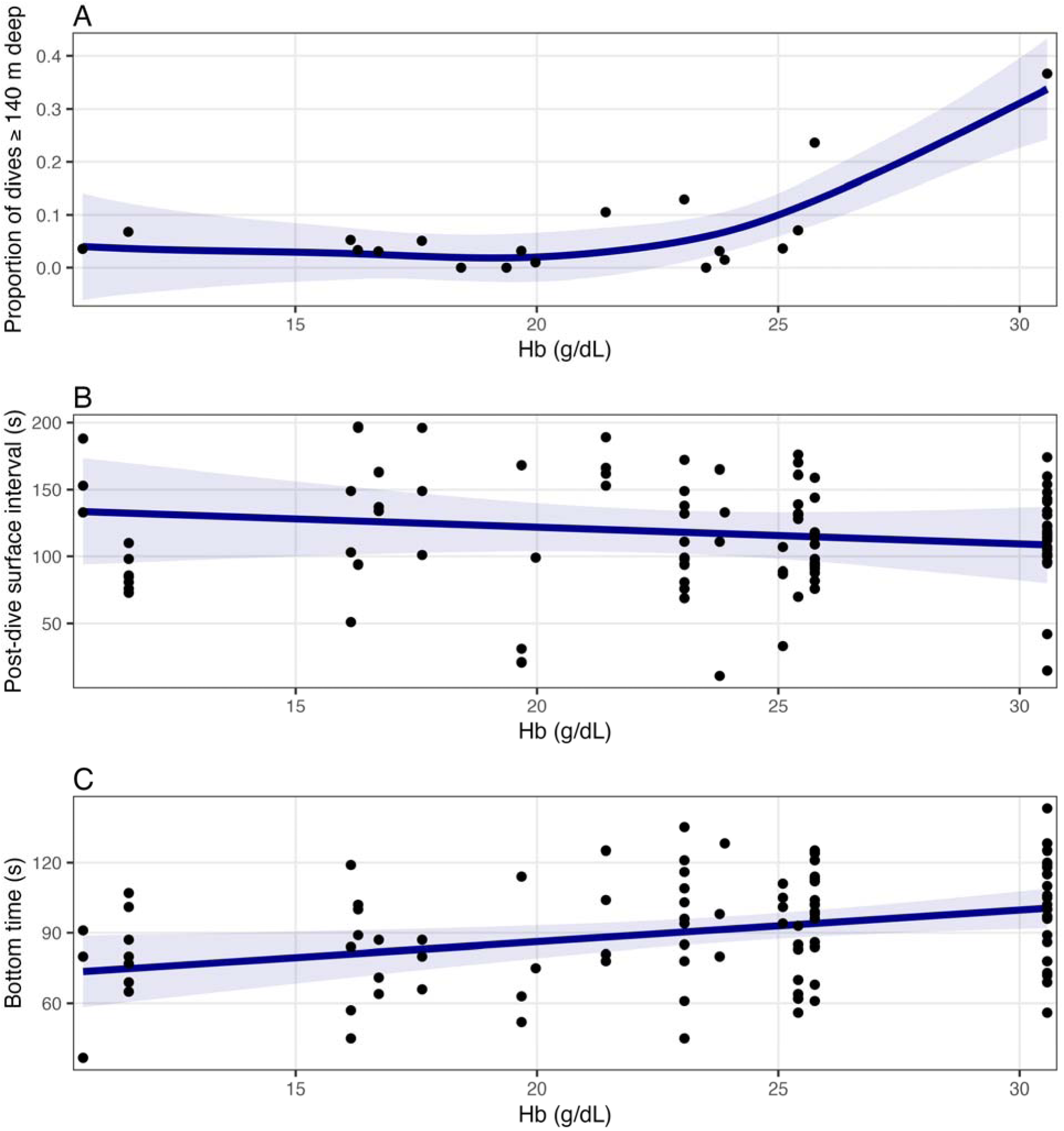
Association between Hb and the proportion of dives an individual made at deeper depth (≥ 140 m, panel A), as well as changes in deep dives’ post-dive surface intervals (panel B) and bottom times (panel C). A total of 19 penguins were included in the analysis from panel A (5 from Race Point and 14 from Pebble Island), but only 16 were included in panels B and C because three individuals did not make dives ≥ 140 m. The blue lines represent the predicted value, and the blue bands represent the 95% confidence intervals. The points represent the observed values.

**Table 2.** Results from the final GAMM for diving efficiency. We present the estimate, standard error (SE) of the parametric coefficient (intercept and fixed effect), as well as their associated test statistic, and p-value. We also present the estimated degrees of freedom (EDF), test statistic, and p-value associated with the smooth terms of the GAMM. For model selection, we used the shrinkage method with a double penalty, which allows non-important terms to be shrunk to the null. A total of 19 penguins were included in this analysis (5 from Race Point and 14 from Pebble Island).

| Term | Parametric coefficient |  |  |  | Significance of smooth term |  |  |
| --- | --- | --- | --- | --- | --- | --- | --- |
|  | Estimate | SE | *t* | p-value | EDF | F | p-value |
| Intercept | 0.295 | 0.006 | 47.606 | < 0.001 |  |  |  |
| Colony (Race point) | 0.034 | 0.014 | 2.474 | 0.014 |  |  |  |
| s(Depth) |  |  |  |  | 7.856 | 33.369 | < 0.001 |
| s(Mass) |  |  |  |  | 1.000 | 0.005 | 0.943 |
| s(Hct) |  |  |  |  | 1.000 | 2.302 | 0.129 |
| s(Hb) |  |  |  |  | 1.000 | 0.140 | 0.708 |
| ti(Depth, Mass) |  |  |  |  | 1.000 | 2.613 | 0.106 |
| ti(Depth, Hct) |  |  |  |  | 10.675 | 3.985 | < 0.001 |
| ti(Depth, Hb) |  |  |  |  | 9.683 | 5.110 | < 0.001 |

**Table 3.**
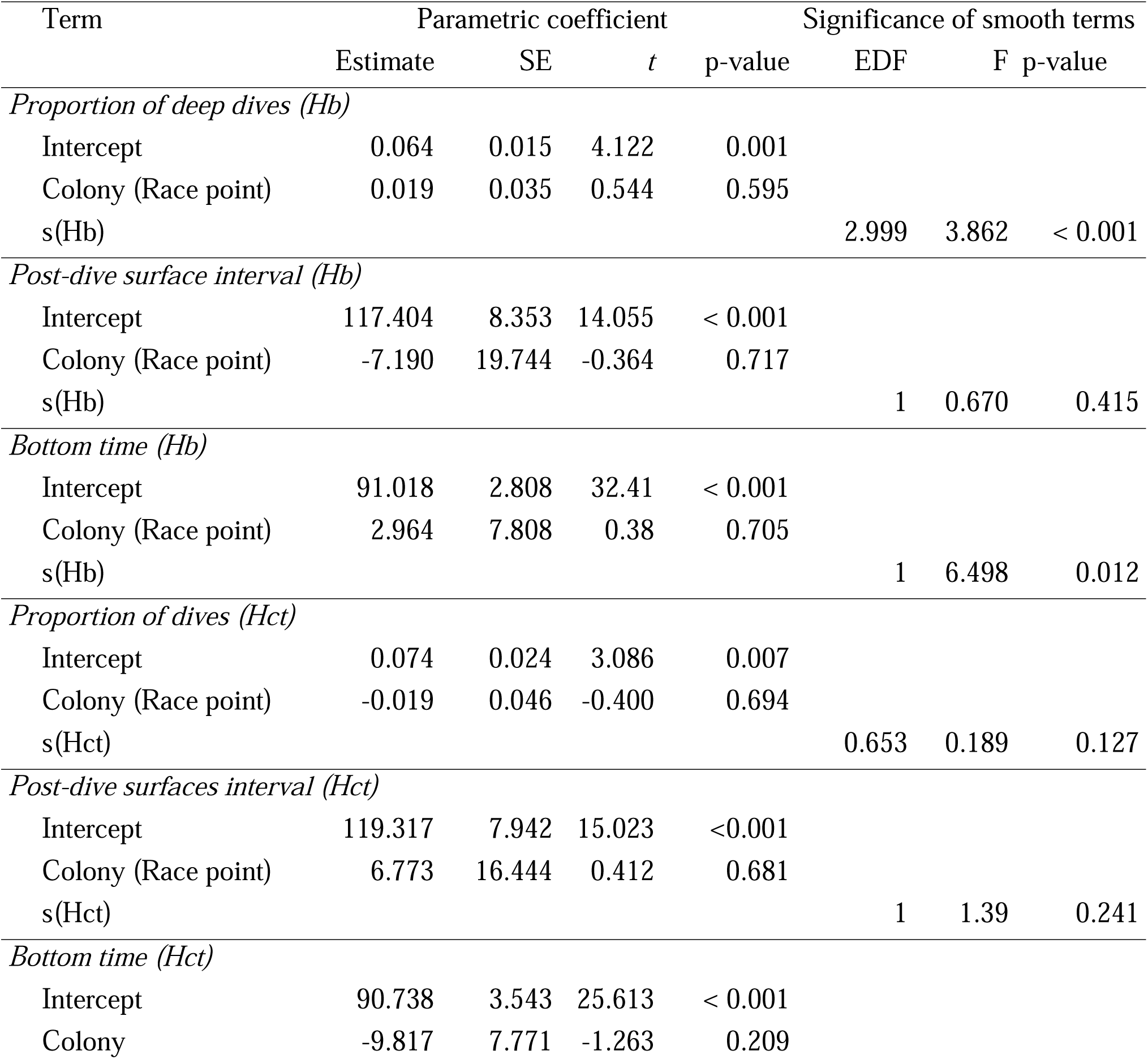
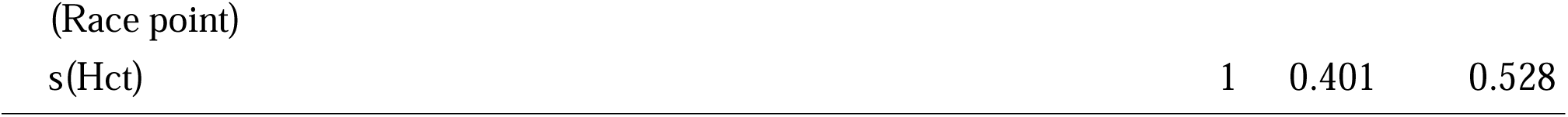
Parameter estimates and test statistic values for the post-hoc GAMs relating the proportion of deep dives to each Hb and Hct, as well as for the post-hoc GAMMs relating either post-dive surface interval of deep dives or bottom time of deep dives to each Hb and Hct. For the proportion of deep dive, 19 penguins were included in the GAMs (5 from Race Point and 14 from Pebble Island). For the post-dive surface interval and bottom time, only 16 were included in the GAMMs because three individuals did not make dives ≥ 140 m.

| Term | Parametric coefficient |  |  |  | Significance of smooth terms |  |  |
| --- | --- | --- | --- | --- | --- | --- | --- |
|  | Estimate | SE | *t* | p-value | EDF | F | p-value |
| *Proportion of deep dives (Hb)* |  |  |  |  |  |  |  |
| Intercept | 0.064 | 0.015 | 4.122 | 0.001 |  |  |  |
| Colony (Race point) | 0.019 | 0.035 | 0.544 | 0.595 |  |  |  |
| s(Hb) |  |  |  |  | 2.999 | 3.862 | < 0.001 |
| *Post-dive surface interval (Hb)* |  |  |  |  |  |  |  |
| Intercept | 117.404 | 8.353 | 14.055 | < 0.001 |  |  |  |
| Colony (Race point) | -7.190 | 19.744 | -0.364 | 0.717 |  |  |  |
| s(Hb) |  |  |  |  | 1 | 0.670 | 0.415 |
| *Bottom time (Hb)* |  |  |  |  |  |  |  |
| Intercept | 91.018 | 2.808 | 32.41 | < 0.001 |  |  |  |
| Colony (Race point) | 2.964 | 7.808 | 0.38 | 0.705 |  |  |  |
| s(Hb) |  |  |  |  | 1 | 6.498 | 0.012 |
| *Proportion of dives (Hct)* |  |  |  |  |  |  |  |
| Intercept | 0.074 | 0.024 | 3.086 | 0.007 |  |  |  |
| Colony (Race point) | -0.019 | 0.046 | -0.400 | 0.694 |  |  |  |
| s(Hct) |  |  |  |  | 0.653 | 0.189 | 0.127 |
| *Post-dive surfaces interval (Hct)* |  |  |  |  |  |  |  |
| Intercept | 119.317 | 7.942 | 15.023 | <0.001 |  |  |  |
| Colony (Race point) | 6.773 | 16.444 | 0.412 | 0.681 |  |  |  |
| s(Hct) |  |  |  |  | 1 | 1.39 | 0.241 |
| *Bottom time (Hct)* |  |  |  |  |  |  |  |
| Intercept | 90.738 | 3.543 | 25.613 | < 0.001 |  |  |  |
| Colony | -9.817 | 7.771 | -1.263 | 0.209 |  |  |  |

The non-linear interaction between maximum depth and Hct was more complex, with multiple and less easily interpretable peaks in diving efficiency across Hct values (Fig. 1C). The post-hoc analyses revealed that neither bottom time, post-dive surface interval, nor the proportion of deep dives showed a statistically significant relationship with Hct (Table 3).

Individuals displayed variation in diving and foraging beyond those described for diving efficiency. For example, the maximum depth reached by individuals during the 40 days ranged from 74.8 to 217.8 m, and trips at sea ranged from less than one day to 24 days. We provide a general description of dive characteristics, foraging effort measures, and oxygen storage and carrying capacity indices in Supplement 5.

### 3.2 Foraging effort relationships with oxygen storage and carrying capacity indices, sex, and reproductive status

Two pre-breeding foraging effort measures were associated with oxygen storage and carrying capacity indices, sex, and reproductive status (Fig. 3, Table 4). The best model for vertical distance travelled indicated that it was increasing with Hb, and the best model for time at sea suggested that early-laying penguins spent less time at sea prior to capture than breeding and non-breeding penguins (Fig 4, Table 4, Supplement 6). However, Table 4 shows that other models had ΔAICc < 4, including the null model for vertical distance travelled. The presence of many competitive models is likely explained by the small sample size (13 individuals) and some of the relationships we observed between covariates. For example, given that the 8 males in our samples had a significantly lower average Hb level than the 5 females (W = 34, p-value = 0.045, n = 13; Fig. 4A), it is not surprising that the second-best model for vertical distance travelled showed that males travelled less than females. Similarly, early-laying penguins had, on average, a lower Hb level than non-breeding penguins (Fig. 4B). Although this difference was not significant (W = 15, p-value = 0.057, n = 8), it may contribute to the fact that the second-best model for time at sea did not include breeding status but included a positive relationship with Hb. Average trip duration had the null model as best model (Table 4), and thus was not considered further. We note that a small sample increases the chance of making a type II error (failing to reject the null hypothesis when it is false) and that AICc strongly penalizes model complexity, thus with a larger sample size more complex relationship may have been detected.

**Figure 3.**
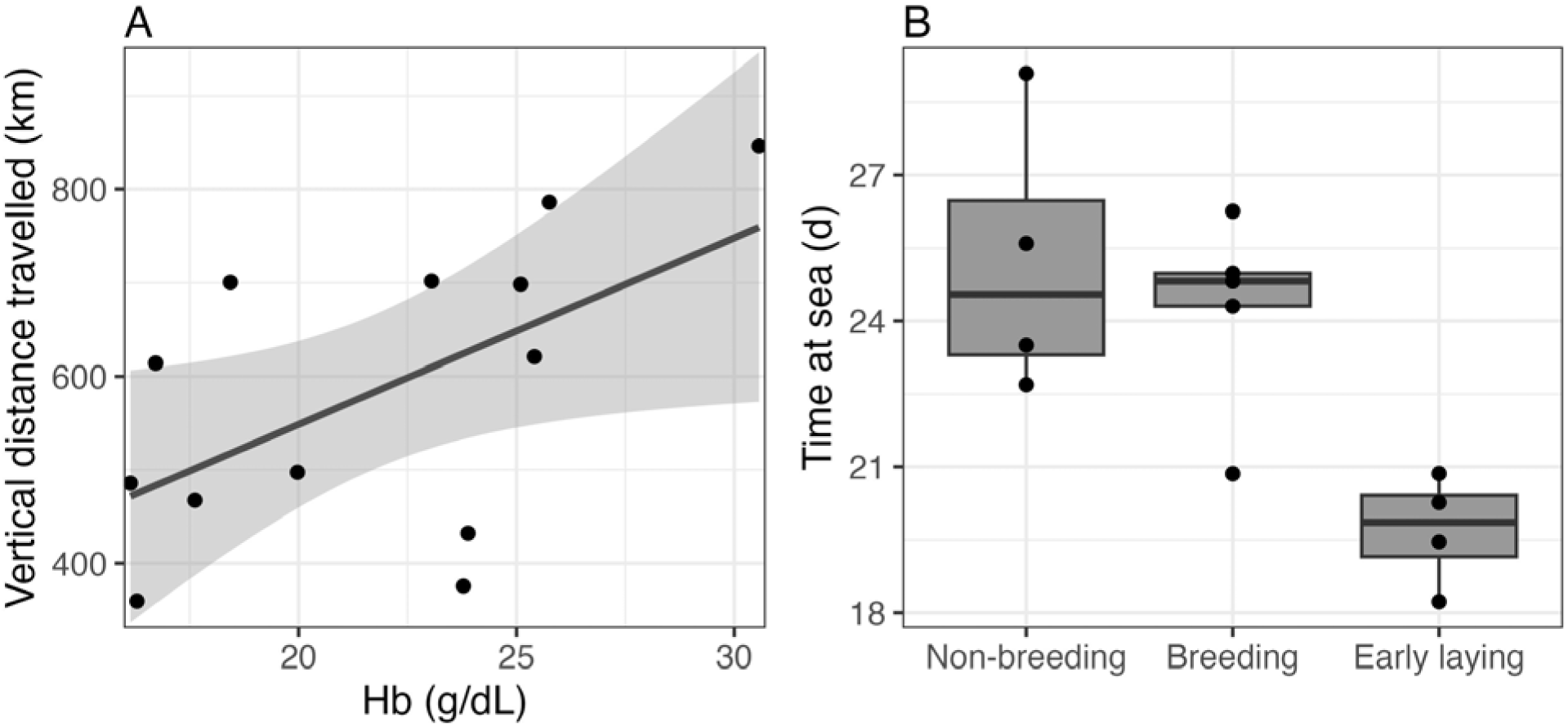
Relationship between Hb and reproductive status and two of the pre-breeding foraging effort measures for the penguins from Pebble Island. Panel A shows the relationship between the sum of the vertical distance travelled while at sea and Hb. The line shows the linear regression line, the band is the 95% confidence interval. Panel B shows the relationship between the time spent at sea and the reproductive status. In both panels, the points are the observations. A total of 13 penguins (4 non-breeding, 5 breeding, and 4 early-laying penguins from Pebble Island) were included in these analyses.

**Figure 4.**
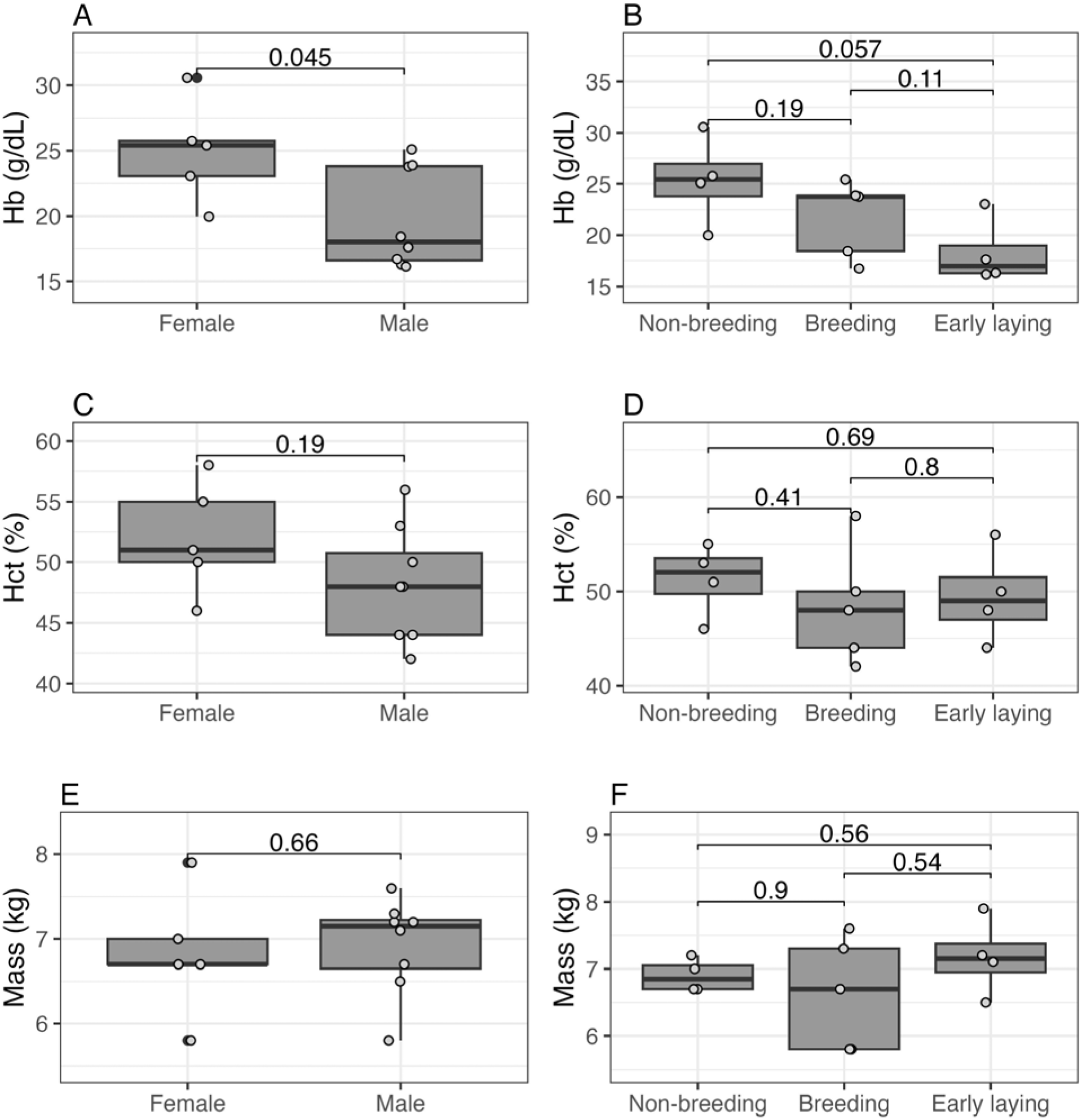
Variation in the three oxygen storage and carrying capacity indices across sex (panels A, C, E) and breeding status (panels B, D, F). Panels A and B show the boxplots (with black points identifying outliers) for Hb, panels C and D for Hct, and panels E and F for mass. In all panels, the grey points are the observations. To be able to see overlapping observation, we added horizontal jitter to the points. The values presented are the p-values associated with the Wilcoxon rank sum tests associated with the pairwise comparisons. A total of 13 penguins (4 non-breeding, 5 breeding, and 4 early-laying penguins from Pebble Island) were included in these analyses.

**Table 4.** Model selection results for the multiple linear regressions relating three measures of foraging effort to oxygen store indices, sex, and breeding status. We present the number of parameters estimated (k), the Akaike Information Criterion with correction for small sample size (AICc), and the difference in AICc from the best model (ΔAICc). While we explore all possible combinations of three covariates for each measure, we only show in the table the null model (no covariates) and models that are either with ΔAICc < 4 of the best model (model with lowest AICc value) or all the models that outperformed the null model if the null model had a ΔAICc ≥ 4. Note that no models with breeding status are included in the table for vertical distance travelled and trip duration, as none of them had a ΔAICc small enough to be part of the subset presented. A total of 13 penguins (all from Pebble Island) were included in these analyses.

| Response | Covariates | k | AICc | $\Delta\text{AICc}$ |
| --- | --- | --- | --- | --- |
| Vertical distance travelled | Hb | 3 | 171.0 | -- |
|  | Sex | 3 | 171.4 | 0.34 |
|  | Hct | 3 | 172.6 | 1.56 |
|  | (Null) | 2 | 172.7 | 1.69 |
|  | Hb + Hct | 4 | 173.8 | 2.76 |
|  | Hb + Sex | 4 | 173.8 | 2.78 |
|  | Hct + Sex | 4 | 173.9 | 2.92 |
| Time at sea | Breeding status | 4 | 66.0 | -- |
|  | Hb + Hct + Mass | 5 | 67.1 | 1.17 |
|  | Hb + Mass | 4 | 67.9 | 1.91 |
|  | Hb | 3 | 68.3 | 2.32 |
|  | Breeding status + Mass | 5 | 69.8 | 3.82 |
|  | Breeding status + Sex | 5 | 70.3 | 4.36 |
|  | Breeding status + Hb | 5 | 70.4 | 4.47 |
|  | (Null) | 2 | 70.5 | 4.51 |
| Trip duration | (Null) | 2 | 59.0 | -- |
|  | Hct | 3 | 59.4 | 0.40 |
|  | Hb + Hct | 4 | 60.4 | 1.48 |
|  | Sex | 3 | 61.8 | 2.82 |
|  | Hb | 3 | 62.0 | 3.00 |
|  | Mass | 3 | 62.3 | 3.39 |
|  | Hct + Mass | 4 | 62.8 | 3.83 |

### 3.3 Potential effects of tagging

Almost all (32/35) tagged penguins recaptured gained weight during the winter, with an average gain of 1.1 kg ± 0.7. However, on average the tagged penguins were significantly lighter at recapture than the untagged penguins (tagged penguins: 6.9 kg ± 0.6, 5.7 - 8.2kg; untagged penguins: 7.3 kg ± 0.5, 6.2 - 8.8 kg; t_66.498_ = 2.812, p-value = 0.006, n = 70). We did not find any other significant differences between tagged and untagged penguins. Their Hb and Hct values were similar (Hb: t_62.863_ = -0.524, p-value = 0.602, n = 66; Hct: t_63.633_ = 1.480, p-value = 0.144, n = 66), and their breeding status did not differ significantly (χ^2^ = 1.167, simulated p-value = 0.604, n = 39). However, given the small sample size for the breeding status analysis (tagged =18, untagged = 19), we note that, compared to the untagged penguins, there was a higher percentage of tagged penguins that did not participate in breeding and a lower percentage of tagged penguins that laid early (Supplement 7). Overall, our results suggest that it is unlikely the relationships we found with blood oxygen storage and carrying capacity indices (i.e., Hb and Hct) are affected by tagging, but that tagging is associated with reduced mass and that the effects of tagging on breeding behaviour and reproductive success should be further explored. See Supplement 7 for more details.

## 4. Discussion

Our results show that the diving efficiency of gentoo penguin was related to Hb, with the efficiency of deep dives (≥ 140 m) increasing with Hb level. This increase in efficiency can be explained by the fact that Hb limits the total amount of oxygen able to be carried by the blood (Minias 2015, Roncon et al. 2018). Of the factors that could have affected diving efficiency, higher Hb was primarily associated with an increase in the bottom time of the dive rather than reduced recovery time (i.e., no decrease in post-dive surface interval with increased Hb levels was observed). Gains in diving efficiency, and associated increased time at the bottom, are likely important to gentoo penguins, as they increase the time available to search for and capture prey during foraging dives. As prey are almost always encountered during the bottom phase of dives (Simeone & Wilson 2003, Takahashi et al. 2004, 2008, Kokubun et al. 2010), increased bottom time translates into time spent where prey is most likely to be encountered. For dives ≥ 140 m, penguins with high Hb (20-31 g/dL) increased their mean bottom phase by 22 % compared with penguins with low Hb (10-20 g/dL). Such an increase could impact foraging success.

The proportion of deep dives performed by penguins increased with Hb levels, with only individuals with high Hb levels regularly diving 140 m or deeper. Vertical distance travelled pre-breeding also increased with Hb levels, suggesting that individuals with high Hb levels may use a deep diving strategy that consequently requires more diving both because they tend to dive deeper and because they have to travel longer in order to reach deep waters further from shore. Such individuals are accessing resources only available at deep depths, giving them access to a wider variety of prey items and potentially decreasing intraspecific competition (e.g., Cimino et al. 2016). Similar findings have been documented in other species. For example, macaroni penguins (*Eudyptes chrysolophus*) forage in habitats within their aerobic capacity (Crossin et al. 2015), and Hb is correlated with higher-quality diets (reviewed in Minias 2015).

Diving efficiency displayed a complex non-linear relationship with Hct levels. The complexity of this non-linear relationship could be associated with the counteracting effects of high Hct levels. While a high Hct level is associated with a higher number of oxygen-carrying red blood cells, it is also linked to increased blood viscosity, and thus increased cardiac workload or decreased blood flow (Birchard 1997, Schuler et al. 2010, Reinhart 2016). In addition, optimum Hct levels can be affected by factors such as age, sex, breeding status, and disease (Reinhart 2016, Johnstone et al. 2017, Brown et al. 2021). In research employing Hct as an index of body condition, higher values are often assumed to indicate better physiological performance (reviewed in Fair et al. 2007, Minias 2015). In contrast, our results suggest that Hct has a complex non-linear relationship with diving efficiency and that higher Hct may not always result in higher diving efficiency. The use of Hct as a physiological indicator has been criticized due to inconsistent relationships with performance and fitness-related traits in many species (reviewed in Fair et al. 2007, Minias 2015, Johnstone et al. 2017). For example, intermediate, rather than high Hct levels were linked to maximum longevity and recruitment rate in house wrens (*Troglodytes aedon*, Bowers et al. 2014). Our results suggest that, while Hct relates to diving efficiency, linear relationships are likely too simplistic. Instead, non-linear models should be used to capture the complex relationship between these variables, though such models require a sufficiently large sample size.

While values of both Hb and Hct aligned with previously reported values for gentoo penguins (20.8 ± 5.4 g/dL and 49.2 ± 4.7 % in our study, n = 66,; 20.7 ± 1.6 g/dL and 50 ± 4 % in Ibañez et al. 2015), they were not closely correlated to one another (see Supplements 2-4). Because mature red blood cells have similar amounts of Hb, we expected a correlation between Hct and Hb. However, ecological stressors can affect these indices differently (Bańbura et al. 2007, Wagner et al. 2008, Johnstone et al. 2017), and weak correlations have been documented in other seabird species (e.g., Kaliński et al. 2011). While some anaemic birds with low Hb concentrations may be unable to improve their condition, some will actively regenerate red blood cells (Jaensch & Clark 2004, Fair et al. 2007, Campbell et al. 2010). In such cases, whether Hb and Hct will remain correlated will be affected by erythropoiesis. Erythropoiesis increases the number of immature red blood cells, which are larger and initially have lower Hb content, disproportionately increasing Hct but not Hb (Wagner et al. 2008, Campbell et al. 2010, Johnstone et al. 2017). We identified one individual that may have demonstrated evidence of compromised health. This non-breeding penguin had the lowest Hb and yet the fourth highest Hct. Anaemia can be associated with additional health conditions, such as heavy parasite loads or severe injury (Vleck et al. 2000, Jaensch & Clark 2004, Fair et al. 2007, Clark 2015). Despite showing no apparent signs of injury at the time of re-capture, this anaemic individual was one of the only individuals that lost mass between April and October. Such an example supports experimental designs that measure both Hb and Hct when assessing individual health (Johnstone et al. 2017).

According to optimal foraging theory, penguins maximize their bottom time and minimize energy spent travelling down to where prey is located (Stephens & Krebs 1986, Wilson et al. 2010, Zimmer et al. 2010). As such, shallow dives should be preferred, requiring less travel time and energy to swim down to depth (Shepard et al. 2009, Zimmer et al. 2010). Deeper dives are only profitable when prey capture rates, size, or caloric value outweigh the energetic costs of diving (Sala et al. 2014b). In our data, peak diving efficiency occurred in shallow foraging habitat at approximately 60 m deep (Fig. 1A), which is deeper than previously reported values (25-30 m; Lee et al. 2015). In line with theory, this putative optimal depth (60 m) was repeatedly visited by penguins (see Supplement 5). However, potentially to reach larger and/or energy-rich prey, many penguins foraged far deeper than this putative optimal depth (maximum depth reached by individuals: 166 m ± 29).

Diet studies suggest that deeper dives may provide greater access to larger prey that may be more valuable to gentoo penguins than the smaller prey in shallower habitat. Clausen et al. (2005) compared the diet of gentoo penguins in colonies around the Falkland Islands with surveys of prey abundance during the 2000 pre-breeding season. They highlight preferential foraging of squid and fish over krill, which has also been highlighted in the historical diet of gentoo penguins (McMahon et al. 2019). Prey likely to be essential for Pebble Island and Race Point penguins includes bentho-demersal lobster krill (*Munida gregaria*), found at depths 26-116 m, and Patagonian squid (*Doryteuthis gahi*), found at depths 60-96 m, both of which were in high abundance close to the colonies in 2000 (Clausen et al. 2005). A deeper squid species (*Moroteuthopsis ingens*), found at depths 100-166 m, and various rock cod species (*Patagontothen spp.*), found at depths 40-205 m, are abundant further offshore and were found in their diet in 2000 (Clausen et al. 2005). These prey species vary in size and energy density. For example, the Patagonian squid found at shallower depth is smaller (max length = 28.0 cm, Palomares & Pauly 2023) but has a slightly higher energy density (5.0 kJ/g; Ciancio et al. 2007) than rock cod species such as *Patagonotothen ramsayi* (max length = 44.4 cm, Froese & Pauly 2023; 4.7 kJ/g, Ciancio et al. 2007). In a more recent assessment of prey, rock cods were found to be a principal prey item for gentoo penguins in the Falkland Islands (Handley et al. 2016). In addition, underwater footage demonstrated that gentoo penguins do not deviate from their path for small prey such as lobster krill, but actively chase larger prey items such as large squid and fish species (Handley et al. 2018). Generally, for marine species, prey size increases with depth (Masello et al. 2010, Miller et al. 2010).

Using diving efficiency as a measure of diving abilities implicitly assumes that animals typically dive to their aerobic dive limit and does not account for the numerous factors influencing dive time and efficiency. For example, predators adapt foraging strategies based on prey conditions (e.g., Magellanic Penguins, *Spheniscus magellanicus*; Sala et al. 2014a) and may abandon a dive early in poor prey conditions (e.g., Thums et al. 2013, Viviant et al. 2016). Such behavioural factors likely explain the low diving efficiency at shallow depths. However, deep diving is more energetically costly to undertake (Shepard et al. 2009, Zimmer et al. 2010). As such, oxygen storage and carrying capacity are more likely to influence the behaviour of animals foraging at deeper depths (Thompson & Fedak 2001, Viviant et al. 2016) and may explain why the relationship with Hb is more apparent at depth.

Gentoo penguins that laid early spent significantly less time at sea than non-breeding penguins, a difference we attribute to higher foraging efficiency rather than higher breeding obligations. Participating in breeding compels individuals to defend nest locations and partake in pair-bonding behaviours, requiring many species of penguins to fast during this time (Williams 1995). While gentoo penguins do not fast during breeding, they increase their time on land to maintain a nest site for approximately two weeks before egg-laying (Black 2016). All monitored gentoo penguins, except one non-breeder, were captured at least 11 days (14.5 ± 2.2 days, September 30 to October 07) before eggs were first seen in the colony (October 18). Thus, while early-laying penguins may have been maintaining nest sites for a few days at the end of the TDR observation period, most of the 40-day data would be prior to nesting and may be associated with early-laying penguins requiring less time to meet energetic demands.

Acquisition of food resources during the pre-breeding period is essential, as breeding birds must expend additional energy to defend their nest and produce eggs (Williams 2012). Successful acquisition of high quality resources relates to earlier laying, which can influence reproductive success (Sorensen et al. 2009). For example, early breeding European shag (*Phalacrocorax aristotelis*) spend less time foraging in winter than other individuals (Daunt et al. 2006), successfully breeding Adélie penguins (*Pygoscelis adeliae*) are more efficient foragers (Lescroël et al. 2010), and breeding Manx shearwater (*Puffinus puffinus*) spend less time in winter foraging and flying than those that defer breeding (Shoji et al. 2015). In species such as the Adélie penguin, non-breeding individuals were 90 % of the mass of breeders (Vleck & Vleck 2002). We found no significant differences in the average mass between non-breeding, breeding, and early-laying penguins. However, in line with what we would expect if early-laying penguins were more efficient foragers, the non-breeding penguins in our sample were 93 % the weight of early-laying penguins. As such, the decrease in time at sea for early-laying gentoo penguins could indicate that these individuals may achieve food requirements in less time during the pre-breeding period.

While none of the indices of oxygen storage and carrying capacity were in the best model for time spent at sea (the best model only included breeding status), Hb, Hct, and mass were included in models with substantial support (ΔAICc < 2), and there are known relationships between these indices and reproduction. In many bird species, Hb levels in females decline during the egg production and laying period (reviewed in Minias 2015). Given the observed increase in diving efficiency with Hb levels, such reduction in Hb could create important trade-offs between foraging capacity and reproduction. Other penguin species, such as macaroni penguins, have evolved extreme size dimorphism due to similar trade-offs between aerobic condition and egg production (Jubinville et al. 2020). As the physiology of each sex is affected differently by breeding (Williams 2012, Desprez et al. 2018), future studies with a larger sample size could explore sex-specific associations between Hb levels, foraging capacity, and breeding, and further disentangle these potential trade-offs.

Foraging is an essential activity for survival, and understanding how an individual’s physiology and health affect dive performance, selection of foraging habitat, and breeding participation is essential to understanding the effects natural and anthropogenic ecosystem changes can have on populations. In the Falkland Islands, the continental shelf is a popular fishing ground, and gentoo penguins are occasionally caught as bycatch and potentially compete for prey with fishing industries (Clausen & Pütz 2003, Trathan et al. 2015). Their extended winter range could overlap with exploration for hydrocarbons, and there is a consideration of further developing fisheries inshore (Augé et al. 2015). Given the associations between an individual’s Hb and Hct levels and their diving and foraging patterns, these human activities could affect portions of the population differently. For example, reducing prey availability nearshore could be especially detrimental for penguins with low oxygen storage and carrying capacity.

According to the island-wide census of 14 colonies, nesting pair numbers and breeding success in 2018 were above the long-term annual average recorded since 2003 (Stanworth and Crofts 2019), suggesting that the penguins in our study experienced relatively prosperous environmental conditions. However, in a year of low environmental conditions, the costs of an inferior strategy could be more prevalent (Fronstin et al. 2016, Storey et al. 2017). Therefore, further understanding the relationship between body conditions and diving efficiency, and its impact on foraging success and breeding participation across a range of environmental conditions, may be crucial in understanding breeding participation and population dynamics in the future.

## Supporting information

Supplement (sections 1-7)

## 5. Acknowledgements

We thank Drs. David Rosen and Colin Brauner for their advice, and Jeff Yap, Mason King, and Elizabeth Ruberg for their assistance with the haemoglobin assays. We thank John and Michelle Jones and Dot and Alex Gould who granted access to the gentoo penguin colonies and provided logistics support. We thank Lina Crossin, Joanna Wong, Katie Harrington, Isabeau Pratte, Megan Boldenow, and Emma Phillips for their assistance in the field, as well as Dr. Sarah Crofts, Dr. Andrew Stanworth, and Riki Evans, along with Falklands Conservation, South Atlantic Environmental Research Institute, and British Antarctic Survey for their logistics support. We thank Dr. Jonathan Handley for his information on tag attachment. Financial support was provided by the National Sciences and Engineering Research Council of Canada (Discovery Grant), the University of British Columbia, and the Werner and Hildegard Hesse Research Award in Ornithology, the Canadian Research Chairs program, Canada Foundation for Innovation (John R. Evans Leaders Fund), and B.C. Knowledge Development Fund.

## 6. Animal care

This research was conducted under Falkland Islands Scientific Research Licence (R12/2017), University of British Columbia Animal Care Permit A17-0243 and Dalhousie University Animal Care Permit 17-100.

## Footnotes

† https://www.accu-metrics.com/animal-services/p/avian-dna-bird-sexing-test

## Notes

### Competing Interest Statement

The authors have declared no competing interest.

### Summary of Updates

Various minor changes throughout to clarify sample size, terms, and results.

## Literature Cited

Alves JA, Gunnarsson TG, Hayhow DB, Appleton GF, Potts PM, Sutherland WJ, Gill JA (2013) Costs, benefits, and fitness consequences of different migratory strategies. Ecology 94:11–17.

Augé A, Lascelles B, Dias MP (2015) Marine spatial planning for the Falkland Islands. ‘Methodology for identification of important areas for marine megafauna’ workshop report. South Atlantic Environmental Research Institute, Stanley, Falkland Islands.

Bańbura J, Bańbura M, Kaliński A, Skwarska J, Słomczyński R, Wawrzyniak J, Zieliński P (2007) Habitat and year-to-year variation in haemoglobin concentration in nestling blue tits Cyanistes caeruleus. Comparative Biochemistry and Physiology, Part A 148:572– 577.

Bannasch R, Wilson RP, Culik B (1994) Hydrodynamic aspects of design and attachment of a back-mounted device in penguins. The Journal of experimental biology 194:83–96.

Baylis AMM, Zuur AF, Brickle P, Pistorius PA (2012) Climate as a driver of population variability in breeding gentoo penguins *Pygoscelis papua* at the Falkland Islands. Ibis 154:30–41.

Baylis AMM, Tierney M, Orben RA, González de la Peña D, Brickle P (2021) Non-breeding movements of gentoo penguins at the Falkland Islands. Ibis 163:507–518.

Bearhop S, Phillips RA, McGill R, Cherel Y, Dawson DA, Croxall JP (2006) Stable isotopes indicate sex-specific and long-term individual foraging specialisation in diving seabirds. Marine Ecology Progress Series 311:157–164.

Beaulieu M, Thierry AM, Handrich Y, Massemin S, le Maho Y, Ancel A (2010) Adverse effects of instrumentation in incubating Adélie Penguins (Pygoscelis adeliae). Polar Biology 33:485–492.

Birchard GF (1997) Optimal hematocrit: theory, regulation and implications. American Zoologist 37:65–72.

Black CE (2016) A comprehensive review of the phenology of Pygoscelis penguins. Polar Biology 39:405–432.

Black C, Collen B, Lunn D, Filby D, Winnard S, Hart T (2018) Time-lapse cameras reveal latitude and season influence breeding phenology durations in penguins. Ecology and Evolution 8: 8286–8296.

Bolnick DI, Svanbäck R, Fordyce JA, Yang LH, Davis JM, Hulsey CD, Forister ML (2003) The ecology of individuals: incidence and implications of individual specialization. The American Naturalist 161:1–28.

Booth JM, Steinfurth A, Fusi M, Cuthbert RJ, McQuaid CD (2018) Foraging plasticity of breeding northern rockhopper penguins, *Eudyptes moseleyi*, in response to changing energy requirements. Polar Biology 41:1815–1826.

Bowers EK, Hodges CJ, Forsman AM, Vogel LA, Masters BS, Johnson BGP, Johnson LS, Thompson CF, Sakaluk SK (2014) Neonatal body condition, immune responsiveness, and hematocrit predict longevity in a wild bird population. Ecology 95:3027–3034.

Brown TJ, Hammers M, Taylor M, Dugdale HL, Komdeur J, Richardson DS (2021) Hematocrit, age, and survival in a wild vertebrate population. Ecology and Evolution 11:214–226.

Burnham KP, Anderson DR (2002) Model selection and multimodel inference: a practical information theoretic approach. 2nd edition. Springer-Verlag New York, Inc., New York, USA.

Campbell TW, Smith SA, Zimmerman KL (2010) Hematology of waterfowl and raptors. In: Weiss DJ, Wardrop KJ (eds) Schalm’s Veterinary Hematology. Sixth edition. Wiley-Blackwell, Ames, USA, p 977–986

Camprasse ECM, Cherel Y, Bustamante P, Arnould JPY, Bost C-A (2017) Intra-and inter-individual variation in the foraging ecology of a generalist subantarctic seabird, the gentoo penguin. Marine Ecology Progress Series 578:227–242.

Carpenter-Kling T, Handley JM, Green DB, Reisinger RR, Makhado AB, Crawford RJM, Pistorius PA (2017) A novel foraging strategy in gentoo penguins breeding at sub-Antarctic Marion Island. Marine Biology 164:1–33.

Ceia FR, Ramos JA (2015) Individual specialization in the foraging and feeding strategies of seabirds: a review. Marine Biology 162:1923–1938.

Chimienti M, Cornulier T, Owen E, Bolton M, Davies IM, Travis JMJ, Scott BE (2017) Taking movement data to new depths: inferring prey availability and patch profitability from seabird foraging behavior. Ecology and Evolution 7:10252–10265.

Ciancio JE, Pascual MA, Beauchamp DA (2007) Energy density of Patagonian aquatic organisms and empirical predictions based on water content. Transactions of the American Fisheries Society 136:1415–1422.

Cimino MA, Moline MA, Fraser WR, Patterson-Fraser DL, Oliver MJ (2016) Climate-driven sympatry may not lead to foraging competition between congeneric top-predators. Scientific Reports 6:18820.

Clark P (2015) Observed variation in the heterophil to lymphocyte ratio values of birds undergoing investigation of health status. Comparative Clinical Pathology 24:1151– 1157.

Clausen AP, Pütz K (2003) Winter diet and foraging range of gentoo penguins (*Pygoscelis papua*) from Kidney Cove, Falkland Islands. Polar Biology 26:32–40.

Clausen AP, Arkhipkin AI, Laptikhovsky VV, Huin N (2005) What is out there: diversity in feeding of gentoo penguins (*Pygoscelis papua*) around the Falkland Islands (Southwest Atlantic). Polar Biology 28:653–662.

Cook TR, Lescroël A, Cherel Y, Kato A, Bost CA (2013) Can foraging ecology drive the evolution of body size in a diving endotherm? PLoS ONE 8:e56297.

Crossin GT, Williams TD (2016) Migratory life histories explain the extreme egg-size dimorphism of Eudyptes penguins. Proceedings of the Royal Society B 283:20161413.

Crossin GT, Phillips RA, Wynne-Edwards KE, Williams TD (2013) Postmigratory body condition and ovarian steroid production predict breeding decisions by female gray-headed albatrosses. Physiological and Biochemical Zoology: Ecological and Evolutionary Approaches 86:761–768.

Crossin GT, Takahashi A, Sakamoto KQ, Trathan PN, Williams TD (2015) Habitat selection by foraging macaroni penguins correlates with hematocrit, an index of aerobic condition. Marine Ecology Progress Series 530:163–176.

Cury PM, Boyd IL, Bonhommeau S, Anker-Nilssen T, Crawford RJM, Furness RW, Mills JA, Murphy EJ, Österblom H, Paleczny M, Piatt JF, Roux J-P, Shannon L, Sydeman WJ (2011) Global seabird response to forage fish depletion - one-third for the birds. Science 334:1703–1707.

Daunt F, Afanasyev V, Silk JRD, Wanless S (2006) Extrinsic and intrinsic determinants of winter foraging and breeding phenology in a temperate seabird. Behavioral Ecology and Sociobiology 59:381–388.

Desprez M, Jenouvrier S, Barbraud C, Delord K, Weimerskirch H (2018) Linking oceanographic conditions, migratory schedules and foraging behaviour during the non-breeding season to reproductive performance in a long-lived seabird. Functional Ecology 32:2040–2053.

Durell SEALeVdit (2000) Individual feeding specialisation in shorebirds: Population consequences and conservation implications. Biological Reviews 75:503–518.

Drabkin D, Austin H (1932) Spectrophotometric studies I. Spectrophotometric constants for common hemoglobin derivatives in human, dog, and rabbit blood. Journal of biological chemistry 98:719–733.

Fair J, Whitaker S, Pearson B (2007) Sources of variation in haematocrit in birds. Ibis 149:535– 552.

Froese R, Pauly D (2023) FishBase. World Wide Web electronic publication. www.fishbase.org (02/2023).

Fronstin RB, Christians JK, Williams TD (2016) Experimental reduction of haematocrit affects reproductive performance in European starlings. Functional Ecology 30:398–409.

Glazier DS (2005) Beyond the ‘3/4-power law’: variation in the intra-and interspecific scaling of metabolic rate in animals. Biological Review 80:611–662.

Handley JM, Baylis AMM, Brickle P, Pistorius P (2016) Temporal variation in the diet of gentoo penguins at the Falkland Islands. Polar Biology 39:283–296.

Handley JM, Connan M, Baylis AMM, Brickle P, Pistorius P (2017) Jack of all prey, master of some: influence of habitat on the feeding ecology of a diving marine predator. Marine Biology 164:82.

Handley JM, Thiebault A, Stanworth A, Schutt D, Pistorius P (2018) Behaviourally mediated predation avoidance in penguin prey: in situ evidence from animal-borne camera loggers. Royal Society Open Science 5:171449.

Hassrick JL, Crocker DE, Teutschel NM, McDonald BI, Robinson RW, Simmons SE, Costa DR (2010) Condition and mass impact oxygen stores and dive duration in adult female northern elephant seals. Journal of Experimental Biology 213:585–592.

Herman RW, Valls FCL, Hart T, Petry MV, Trivelpiece WZ, Polito MJ (2017) Seasonal consistency and individual variation in foraging strategies differ among and within Pygoscelis penguin species in the Antarctic Peninsula region. Marine Biology 164:1–13.

Hope ACA (1968) A simplified Monte Carlo significance test procedure. Journal of the Royal Statistical Society Series B 30:582–598.

Horswill C, Trathan PN, Ratcliffe N (2017) Linking extreme interannual changes in prey availability to foraging behaviour and breeding investment in a marine predator, the macaroni penguin. PLoS ONE 12:e0184114.

Hudson LN, Isaac NJB, Reuman DC (2013) The relationship between body mass and field metabolic rate among individual birds and mammals. Journal of Animal Ecology 82:1009–1020.

Ibañez AE, Najle R, Larsen K, Pari M, Figueroa A, Montalti D (2015) Haematological values of three Antarctic penguins: Gentoo (*Pygoscelis papua*), Adélie (*P. adeliae*) and chinstrap (*P. antarcticus*). Polar Research 34:25718.

Jaensch S, Clark P (2004) Haematological characteristics of response to inflammation or traumatic injury in two species of black cockatoos: *Calyptorhynchus magnificus* and *C. funereus*. Comparative Clinical Pathology 13:9–13.

Johnstone CP, Lill A, Reina RD (2017) Use of erythrocyte indicators of health and condition in vertebrate ecophysiology: a review and appraisal. Biological Reviews 92:150–168.

Jubinville I, Williams TD, Trathan PN, Crossin GT (2020) Trade-off between aerobic performance and egg production in migratory macaroni penguins. Comparative Biochemistry and Physiology-Part A: Molecular and Integrative Physiology 247: 110742

Kaliński A, Markowski M, Bańbura M, Mikus W, Skwarska J, Wawrzyniak J, Gladalski M, Zieliński P, Bańbura J (2011) Weak correlation between haemoglobin concentration and haematocrit of nestling Great Tits *Parus major* and Blue Tits *P. caerule*. Ornis Fennica 88:234–240.

Kokubun N, Takahashi A, Mori Y, Watanabe S, Shin H-C (2010) Comparison of diving behavior and foraging habitat use between chinstrap and gentoo penguins breeding in the South Shetland Islands, Antarctica. Marine Biology 157:811–825.

Labocha MK, Hayes JP (2012) Morphometric indices of body condition in birds: a review. Journal of Ornithology 153:1–22.

Lee WY, Kokubun N, Jung J-W, Chung H, Kim J-H (2015) Diel diving behavior of breeding gentoo penguins on King George Island in Antarctica. Animal Cells and Systems 19:274–281.

Lescroël A, Bost CA (2005) Foraging under contrasting oceanographic conditions: the gentoo penguin at Kerguelen Archipelago. Marine Ecology Progress Series 302:245–261.

Lescroël A, Ballard G, Toniolo V, Barton KJ, Wilson PR, Lyver PO, Ainley DG (2010) Working less to gain more: when breeding quality relates to foraging efficiency. Ecology 91:2044– 2055.

Luque SP, Fried R (2011) Recursive filtering for zero offset correction of diving depth time series with GNU R package diveMove. PLoS ONE 6:e15850.

Marra G, Wood SN (2011) Practical variable selection for generalized additive models. Computational Statistics and Data Analysis 55:2372–2387.

Masello JF, Mundry R, Poisbleau M, Demongin L, Voigt CC, Wikelski M, Quillfeldt P (2010) Diving seabirds share foraging space and time within and among species. Ecosphere 1:1–

McMahon KW, Michelson CI, Hart T, McCarthy MD, Patterson WP, Polito MJ (2019) Divergent trophics responses of sympatric penguin species to historic anthropogenic exploitation and recent climate change. Proceedings of the National Academy of Sciences:201913093.

Miller AK, Kappes MA, Trivelpiece SG, Trivelpiece WZ (2010) Foraging-niche separation of breeding gentoo and chinstrap penguins, South Shetland Islands, Antarctica. The Condor 112:683–695.

Minias P (2015) The use of haemoglobin concentrations to assess physiological condition in birds: a review. Conservation Physiology 3:cov007.

Minias P (2020) Ecology and evolution of blood oxygen-carrying capacity in birds. American Naturalist 195:788–801.

Minias P, Kamiński M, Janiszewski T, Indykiewicz P, Kowalski J, Jakubas D (2023) Experimental reduction in blood oxygen-carrying capacity alters foraging behaviour in a colonial waterbird. Journal of Experimental Biology. 10.1242/JEB.245443

Mirceta S, Signore AV, Burns JM, Cossins AR, Campbell KL, Berenbrink M (2013) Evolution of Mammalian Diving Capacity Traced by Myoglobin Net Surface Charge Evolution of Mammalian Diving. Science 340:1234192.

Nunes, A. 2019. Divebomb, version 1.0.7. Ocean Tracking Network, Halifax, CA.

Otley HM, Clausen AP, Christie DJ, Pütz K (2004) Aspects of the breeding biology of the gentoo penguin *Pygoscelis papua* at volunteer beach, Falkland Islands, 2001/02. Marine Ornithology 32:167–171.

Palomares MLD, Pauly D (2023) SeaLifeBase. World Wide Web electronic publication. www.sealifebase.org, version (04/2023)

Phillips RA, Lewis S, González-Solís J, Daunt F (2017) Causes and consequences of individual variability and specialization in foraging and migration strategies of seabirds. Marine Ecology Progress Series 578:117–150.

Photopoulou T, Fedak MA, Thomas L, Matthiopoulos J (2014) Spatial variation in maximum dive depth in gray seals in relation to foraging. Marine Mammal Science 30:923–938.

Pistorius PA, Huin N, Crofts S (2010) Population change and resilience in gentoo penguins *Pygoscelis papua* at the Falkland Islands. Marine Ornithology 38:49–53.

Polito MJ, Trivelpiece WZ, Patterson WP, Karnovsky NJ, Reiss CS, Emslie SD (2015) Contrasting specialist and generalist patterns facilitate foraging niche partitioning in sympatric populations of Pygoscelis penguins. Marine Ecology Progress Series 519:221– 237.

Ponganis PJ, Kooyman GL (2000) Diving physiology of birds: a history of studies on polar species. Comparative Biochemistry and Physiology, Part A 126:143–151.

Python Software Foundation. 2019. Python Language Reference, version 3.4.10. Amsterdam, AN.

R Core Team (2022) R: A language and environment for statistical computing. R Foundation for Statistical Computing, Vienna, Austria. https://www.R-project.org/

Reinhart WH (2016) The optimum hematocrit. Clinical Hemorheology and Microcirculation 64:575–585.

Rodnan GP, Ebaugh FGJ, Fox MRS, Chambers DM (1957) The life span of the red blood cell and the red blood cell volume in the chicken, pigeon and duck as estimated by the use of Na_2_Cr^51^O_4_: with observations on red cell turnover rate in the mammal, bird and reptile. Blood 12:355–366.

Roncon G, Bestley S, McMahon CR, Wienecke B, Hindell MA (2018) View from below: inferring behavior and physiology of Southern Ocean marine predators from dive telemetry. Frontiers in Marine Science 5:464.

Sala JE, Wilson RP, Frere E, Quintana F (2014a) Flexible foraging for finding fish: variable diving patterns in magellanic penguins *Spheniscus magellanicus* from different colonies. Journal of Ornithology 155:801–817.

Sala JE, Wilson RP, Quintana F (2014b) Foraging effort in magellanic penguins: balancing the energy books for survival? Marine Biology 162:501–514.

Schamber JL, Esler D, Flint PL (2009) Evaluating the validity of using unverified indices of body condition. Journal of Avian Biology 40:49–56.

Schuler B, Arras M, Keller S, Rettich A, Lundby C, Vogel J, Gassmann M (2010) Optimal hematocrit for maximal exercise performance in acute and chronic erythropoietin-treated mice. Proceedings of the National Academy of Sciences of the United States of America 107:419–423.

Shaw AK (2020) Causes and consequences of individual variation in animal movement. Movement Ecology 8:1–12.

Shepard ELC, Wilson RP, Quintana F, Laich AG, Forman DW (2009) Pushed for time or saving on fuel: fine-scale energy budgets shed light on currencies in a diving bird. Proceedings of the Royal Society B 276:3149–3155.

Shoji A, Aris-Brosou S, Culina A, Fayet A, Kirk H, Padget O, Juarez-Martinez I, Boyle D, Nakata T, Perrins CM, Guilford T (2015) Breeding phenology and winter activity predict subsequent breeding success in a trans-global migratory seabird. Biology Letters 11:20150671.

Simeone A, Wilson RP (2003) In-depth studies of magellanic penguin (*Spheniscus magellanicus*) foraging: can we estimate prey consumption by perturbations in the dive profile? Marine Biology 143:825–831.

Sorensen MC, Hipfner JM, Kyser TK, Norris DR (2009) Carry-over effects in a Pacific seabird: stable isotope evidence that pre-breeding diet quality influences reproductive success. Journal of Animal Ecology 78:460–467.

Stanworth A, Crofts S (2019) Falkland Islands seabird monitoring programme annual report 2018/2019 (SMP26). Falklands Conservation, Stanley.

Stephens DW, Krebs JR (1986) Foraging theory. Princeton University Press, Princeton, NJ.

Storey AE, Ryan MG, Fitzsimmons MG, Kouwenberg A, Takahashi LS, Robertson GJ, Wilhelm SI, Mckay DW, Herzberg GR, Mowbray FK, Macmillan L, Walsh CJ (2017) Balancing personal maintenance with parental investment in a chick-rearing seabird: physiological indicators change with foraging conditions. Conservation Physiology 5:cox055.

Takahashi A, Dunn MJ, Trathan PN, Croxall JP, Wilson RP, Sato K, Naito Y (2004) Krill-feeding behaviour in a chinstrap penguin, *Pygoscelis antarctica*, compared with fish-eating in magellanic penguins *Spheniscus megellanicus*: a pilot study. Marine Ornithology 32:47–54.

Takahashi A, Kokubun N, Mori Y, Shin HC (2008) Krill-feeding behaviour of gentoo penguins as shown by animal-borne camera loggers. Polar Biology 31:1291–1294.

Tanton JL, Reid K, Croxall JP, Trathan PN (2004) Winter distribution and behaviour of gentoo penguins *Pygoscelis papua* at South Georgia. Polar Biology 27:299–303.

Taylor SS, Leonard ML, Boness DJ, Majluf P (2001) Foraging trip duration increases for Humboldt Penguins tagged with recording devices. Journal of Avian Biology 32:369–372.

Thiebot J, Lescroël A, Pinaud D, Trathan PN, Bost C-A (2011) Larger foraging range but similar habitat selection in non-breeding versus breeding sub-Antarctic penguins. Antarctic Science 23:117–126.

Thompson D, Fedak MA (2001) How long should a dive last? A simple model of foraging decisions by breath-hold divers in a patchy environment. Animal Behaviour 61:287–296.

Thums M, Bradshaw CJA, Sumner MD, Horsburgh JM, Hindell MA (2013) Depletion of deep marine food patches forces divers to give up early. Journal of Animal Ecology 82:72–83.

Trathan PN, García-Borboroglu P, Boersma D, Bost C-A, Crawford RJM, Crossin GT, Cuthbert RJ, Dann P, Davis LS, De La Puente S, Ellenberg U, Lynch HJ, Mattern T, Pütz K, Seddon PJ, Trivelpiece W, Wienecke B (2015) Pollution, habitat loss, fishing, and climate change as critical threats to penguins. Conservation Biology 29:31–41.

van der Hoop JM, Fahlman A, Hurst T, Rocho-Levine J, Shorter KA, Petrov V, Moore MJ (2014) Bottlenose dolphins modify behavior to reduce metabolic effect of tag attachment. The Journal of Experimental Biology 217:4229–4236.

Vandenabeele SP, Shepard ELC, Grémillet D, Butler PJ, Martin GR, Wilson RP (2015) Are bio-telemetric devices a drag? effects of external tags on the diving behaviour of great cormorants. Marine Ecology Progress Series 519:239–249.

Viviant M, Jeanniard-du-Dot T, Monestiez P, Authier M, Guinet C (2016) Bottom time does not always predict prey encounter rate in Antarctic fur seals. Functional Ecology 30:1834– 1844.

Vleck CM, Vleck D (2002) Physiological condition and reproductive consequences in adélie penguins. Integrative and Comparative Biology 42:76–83.

Vleck CM, Vertalino N, Vleck D, Bucher TL (2000) Stress, corticosterone, and heterophil to lymphocyte ratios in free-living adélie penguins. The Condor 102:392–400.

Wagner EC, Stables CA, Williams TD (2008) Hematological changes associated with egg production: direct evidence for changes in erythropoiesis but a lack of resource dependence? The Journal of Experimental Biology 211:2960–2968.

Williams TD (1995) The penguins: Spheniscidae. J. N. Davies, editor. Oxford University Press.

Williams TD (2012) Physiological adaptations for breeding in birds. Princeton University Press.

Wilson RP, Grant WS, Duffy DC (1986) Recording devices on free-ranging marine animals: does measurement affect foraging. Ecology 67:1091–1093.

Wilson RP, Shepard ELC, Laich AG, Frere E, Quintana F (2010) Pedalling downhill and freewheeling up; a penguin perspective on foraging. Aquatic Biology 8:193–202.

Wilson RP, Sala JE, Gómez-Laich A, Ciancio J, Quintana F (2015) Pushed to the limit: food abundance determines tag-induced harm in penguins. Animal Welfare 24:37–44.

Wood SN (2011) Fast stable restricted maximum likelihood and marginal likelihood estimation of semiparametric generalized linear models. Journal of the Royal Statistical Society (B) 73:3–36.

Zimmer I, Wilson RP, Beaulieu M, Ropert-Coudert Y, Kato A, Ancel A, Plötz J (2010) Dive efficiency versus depth in foraging emperor penguins. Aquatic Biology 8:269–277.

