## Supplement (sections 1-7) for "Diving efficiency at depth and pre-breeding foraging effort increase with haemoglobin levels in gentoo penguins"

**Marine Ecology Progress Series**

^1^ Institute for the Oceans and Fisheries, University of British Columbia, Vancouver, British Columbia, Canada, V6T 1Z4

^2^ Department of Biology, Dalhousie University, Halifax, Nova Scotia, B3H 4R2

^3^ Joint Nature Conservation Committee, Quay House, 2 Station Road, Fletton Quays, Peterborough, PE2 8YY, UK.

^4^ South Atlantic Environmental Research Institute, Stanley Cottage, Ross Road, Stanley, FIQQ 1ZZ, Falkland Islands

^5^ School of Biological Sciences (Zoology), University of Aberdeen, Tillydrone Avenue, Aberdeen, Scotland AB24 2TZ, UK

^6^ British Antarctic Survey, High Cross, Madingley Road, Cambridge CB3 0ET UK

^7^ National Oceanography Centre, Waterfront Campus European Way, SO14 3ZH UK

^8^ Department of Biological Sciences, Simon Fraser University, Burnaby, British Columbia, Canada, V5A 1S6

^9^ Department of Statistics, University of British Columbia, Vancouver, British Columbia, Canada, V6T 1Z4

**Supplement 1. Additional information on tag attachment methods**

We equipped each penguin with a Sirtrack K2G 173A SWS KIWISAT 202B Argos tag (34g, 55 x 27 x17 mm, Sirtrack, Havelock North, New Zealand) and a Lotek LAT1800 TDR tag (13.6 g, 62 x 13 x 13 mm, Lotek Wireless Inc, St. John’s, Canada). As in Handley et al. (2018) and Baylis et al. (2021), we attached the devices to midline back feathers using overlapping layers of TESA^®^ tape (Beiersdorf AG, GmbH, Hamburg, Germany) and sealed the tape seams with cyanoacrylate glue (Fig. S1 A-B). To limit extensive feather damage, we did not attach the tags to the feathers directly with epoxy or cyanoacrylate glue (Wilson et al. 1997, Houstin et al. 2022). In fact, the penguins resighted with their tag missing had feathers in relatively good condition (Fig. S1 C-D). Given how many tags were lost and the evidence of poorly affixed tags at recaptured (Fig. S1 C), we would suggest using alternative methods in the future. For example, one could explore the method that Houstin et al. (2022) described for emperor penguins, which combines TESA^®^ tape, cyanoacrylate glue, polyamide cable ties, and epoxy glue on the mounting (not feathers). We do not know why all but one Argos tag fell, while many TDR tags stayed attached. We suggest further studies on the placement, shape, and size of tags, first looking at attaching the tag lower on the back to limit access and potential drag.

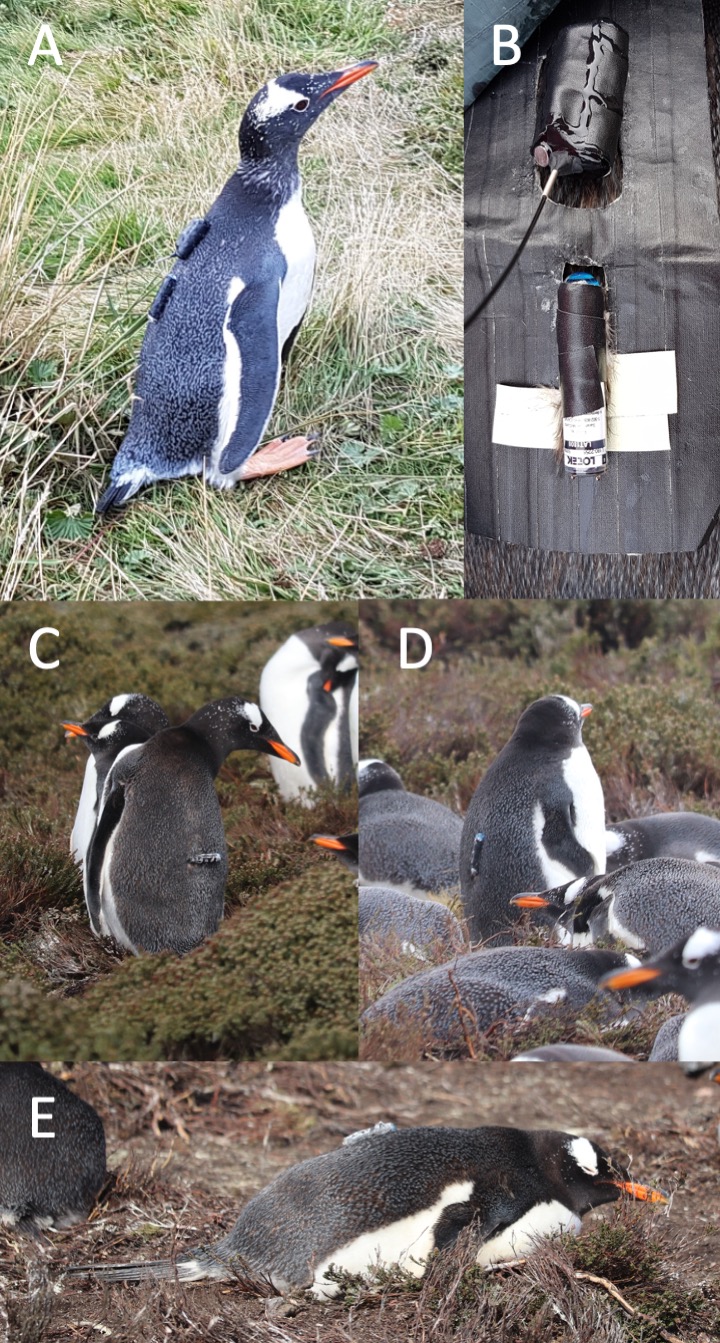
 **Figure S1**. Photographs of tags on penguins at deployment (panels A-B) and recapture (panels C-E). Panel A shows an individual just after the tag was attached. We see the Argos tag at the top of the back with its antennae and the TDR tag just below. Panel B shows the cyanoacrylate glue drying on the seams of the tape holding the Argos tag to the back feather and the tape partly placed on the TDR tag. It shows the stencil we placed on the feathers when we deployed the tag. The stencil was used to make sure all tags were placed at the same distance from one another, that only the good feathers were attached to the tags, and that glue did not fall on other feathers. The stencil was removed once the tags were attached to the penguin. Panel C shows a TDR tag that is no longer fully attached. It was removed subsequently. Panels D and E show TDR tags still well attached but with signs of wear. They were removed subsequently. Panels C and D show that the feathers in the areas where the Argos tags were originally placed appear in relatively good condition.

**Supplement 2. Assessing whether sex should be included in the analysis of dive efficiency**

Given that we had three individuals for whom we did not have sex data, and a small dataset, we wanted to assess whether we could ignore sex differences so as to increase our sample size. To do so, we applied a Generalized additive mixed model (GAMM) to the 16 individuals for which we had sex information. As described in the main text, we first thinned the sample to remove temporal autocorrelation (kept one dive every 70 dives). We then used the R package mgcv (Wood 2011) to apply our model to our data. Our response variable was the efficiency of a dive. We included sex and colony as fixed effects, and penguin id as a random effect. As in the main text, to explore the relationship between dive efficiency, maximum depth of a dive, and the indices of oxygen storage and carrying capacity, we included thin-plate regression splines for maximum depth, Hb, Hct, and mass, as well as tensor product terms (cubic regression splines) for the interactions between maximum depth and each of the oxygen storage and indices. We used the double penalty method described in Marra & Wood (2011) to penalize unimportant covariates. In addition, we applied another GAMM to the same sample that did not include sex, and used Akaike Information Criterion with correction for small sample size (AICc) to compare the models with and without sex.

The GAMM model estimates and test statistic value indicated that sex was not an important covariate (Table S1). In addition, the AICc value of the model without sex (fit to the same dataset) was lower than that of the model with sex (ΔAICc = 2.049). Overall, the results from the model with sex were very similar to the results presented in the main text. Importantly, all the same variables appear important (Table S1). Second, while there are some numerical differences, the general patterns remain the same (Table S1, Fig. S2). The only small notable difference is that the interaction between maximum depth and Hb appears more linear when sex is included (Table S1), and this may be due to the fact that Hb values differ between sexes (Fig. 4 in main text). However, the main interpretation remains the same: higher Hb values are associated with increased dive efficiency at depth.

**Table S1.** Parameter estimates and test statistic values for the GAMM for dive efficiency that includes sex as a covariate. A total of 16 penguins were included in this analysis.

| Term | Parametric coefficient | | | | Significance of smooth terms | | |
| --- | --- | --- | --- | --- | --- | --- | --- |
|  | Estimate | SE | *t* | p-value | EDF | F | p-value |
| Intercept | 0.296 | 0.012 | 25.236 | < 0.001 |  |  |  |
| Colony (Race point) | 0.039 | 0.016 | 2.409 | 0.016 |  |  |  |
| Sex (Male) | -0.002 | 0.016 | -0.132 | 0.895 |  |  |  |
| s(Depth) |  |  |  |  | 7.918 | 24.903 | < 0.001 |
| s(Mass) |  |  |  |  | 1.000 | 0.152 | 0.697 |
| s(Hct) |  |  |  |  | 1.000 | 3.501 | 0.062 |
| s(Hb) |  |  |  |  | 1.000 | 0.005 | 0.944 |
| ti(Depth, Mass) |  |  |  |  | 1.000 | 1.286 | 0.257 |
| ti(Depth, Hct) |  |  |  |  | 11.729 | 4.549 | < 0.001 |
| ti(Depth, Hb) |  |  |  |  | 1.000 | 10.591 | 0.001 |

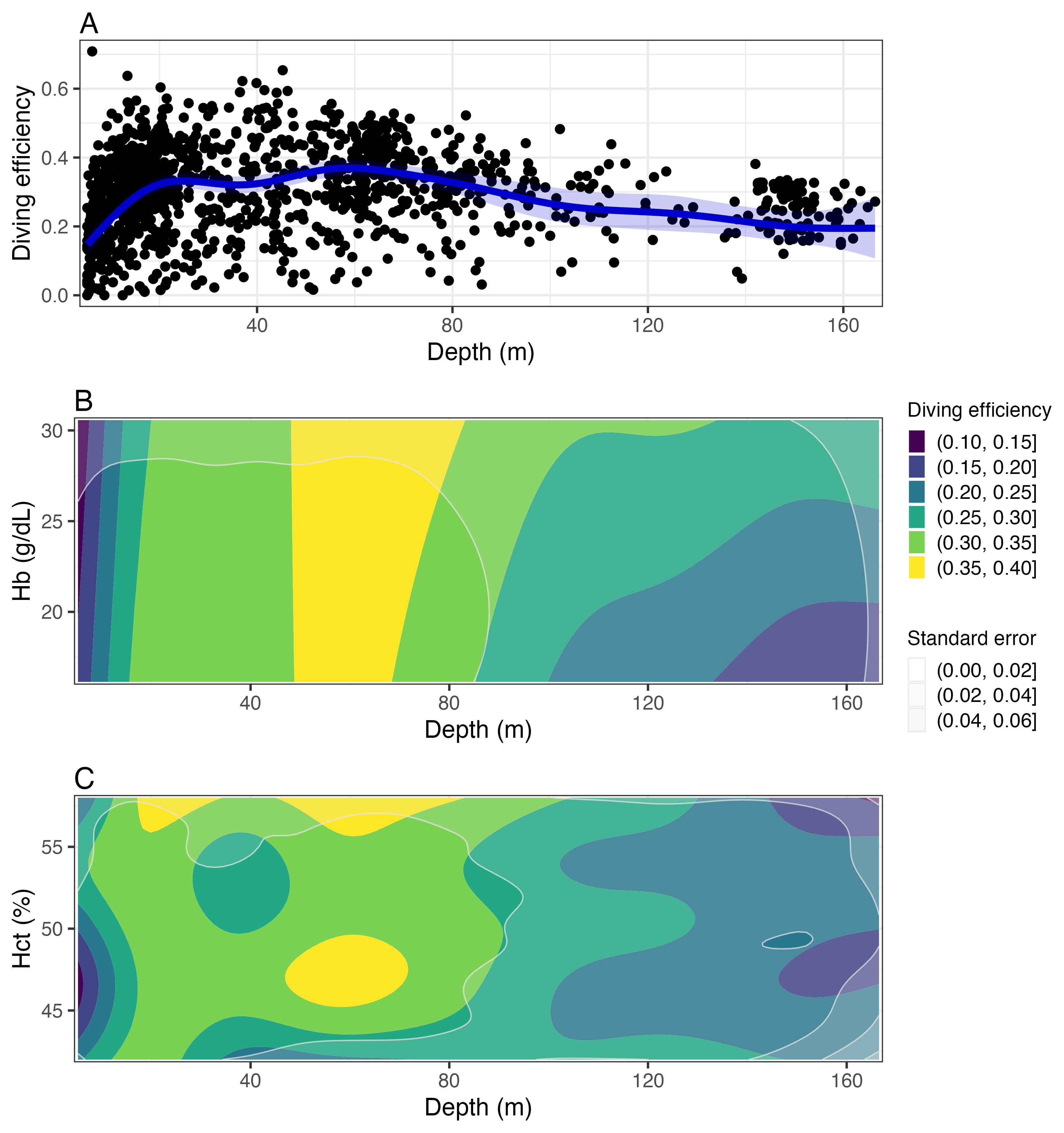

**Figure S2.** Slices of predictions from the GAMM with sex included as a fixed effect. Panel A displays how the predicted dive efficiency changes in response to varying maximum depth values of a dive. The points show the observed values. Panel B shows how dive efficiency is predicted to change with both depth and Hb, while panel C shows how dive efficiency is predicted to change with both depth and Hct. For both these panels, the color represents the dive efficiency value (yellow representing highest efficiency) and the grey bands represent the standard error (more opaque grey representing higher standard error). The areas with larger standard error values (i.e., more opaquely shaded) should be interpreted more cautiously. A total of 16 penguins were included in this analysis.

**Supplement 3. Correlation between covariates and concurvity**

We expect some of the oxygen storage and carrying capacity indices to be correlated, and as such may expect some concurvity to occur in our analysis of dive efficiency. Here, we assess the level of correlation between the three indices of oxygen storage and carrying capacity and the concurvity between the smooth terms of the final GAMM. To see whether the interpretation of the final model is affected by the presence of correlation and concurvity, we also apply simpler GAMMs, which have only one of the oxygen storage and carrying capacity indices deemed important in the final model. We compared the results of these simpler models to the final model presented in the main text.

The three oxygen storage and carrying capacity indices showed marginally important correlation (Fig. S3), with Hct and Hb showing the highest correlation (correlation coefficient of 0.394). In addition, the final GAMM appeared to exhibit important concurvity between its components (Table S2). However, the new GAMMs which included only one of the important oxygen storage and carrying capacity indices at a time displayed similar relationships as those found in the final model presented in the main text (Fig. S4). The dive efficiency increased at medium depths, and then decreased at deeper depth (Fig. S4A-B). As in the final model presented in the main text, the dive efficiency appeared to be mostly increasing with Hb at deeper depths (Fig. S4C) and the dive efficiency had a complex relationship with Hct across depth (Fig. S4D).

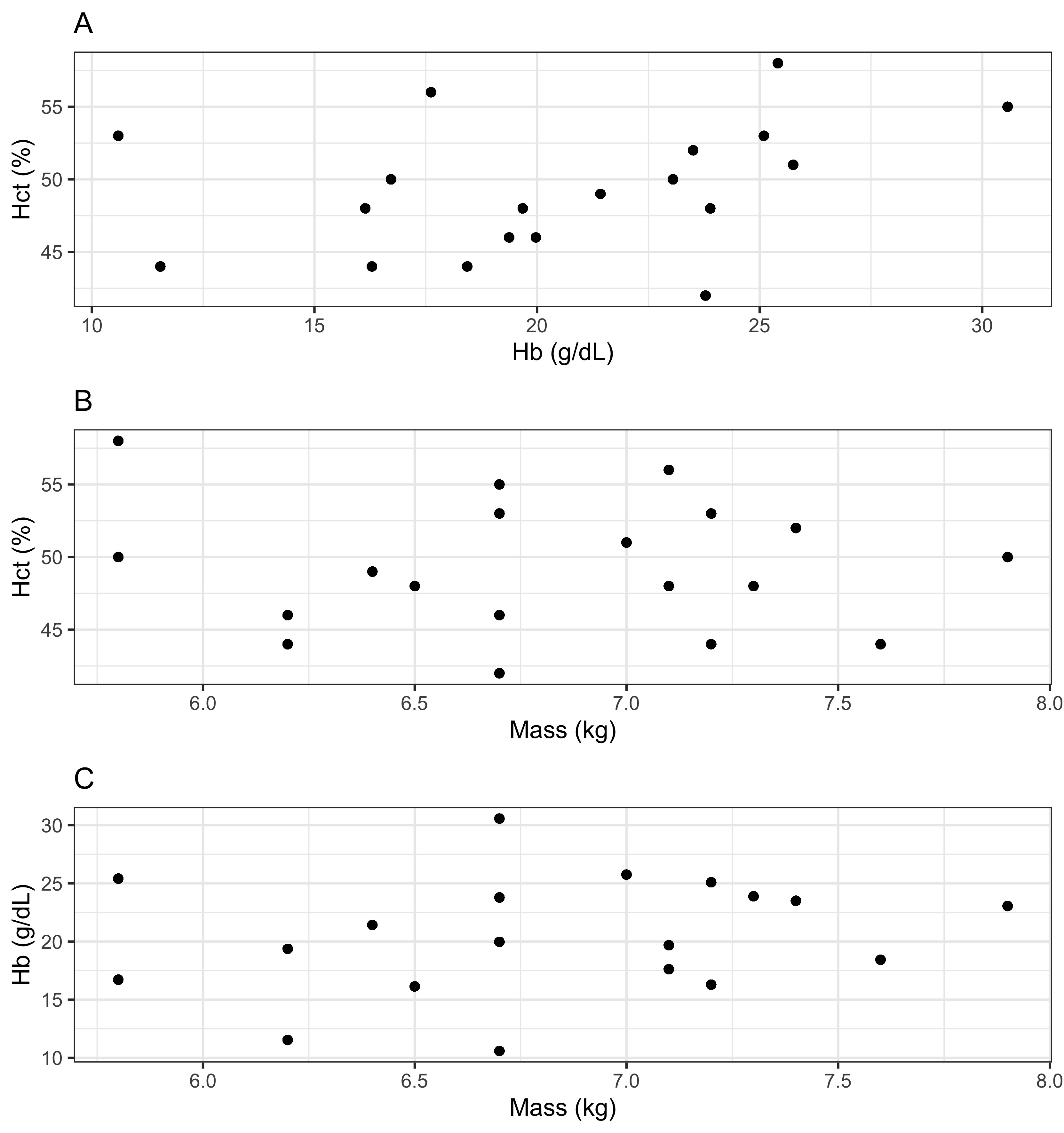

**Figure S3.** Correlation between Hb, Hct, and mass for the 19 penguins used in the final dive efficiency analysis. Panel A shows the Hb and Hct values, which have a correlation coefficient of 0.394. Panel B shows the Mass and Hct values, which have a correlation coefficient of -0.175. Panel C shows the Mass and Hb values, which have a correlation coefficient of 0.180.

**Table S2.** Summary measures of concurvity between the fixed parameters (Para) and the smooth components of the final GAMM, as returned by the mgcv function concurvity.

|  | Para | s(Depth) | s(Mass) | s(Hct) | S(Hb) | ti(Depth, Mass) | ti(Depth, Hct) | ti(Depth, Hb) |
| --- | --- | --- | --- | --- | --- | --- | --- | --- |
| Worst | 1 | 0.680 | 1 | 1 | 1 | 0.911 | 0.951 | 0.934 |
| Observed | 1 | 0.348 | 1 | 1 | 1 | 0.620 | 0.608 | 0.603 |
| Estimate | 1 | 0.630 | 1 | 1 | 1 | 0.546 | 0.623 | 0.569 |

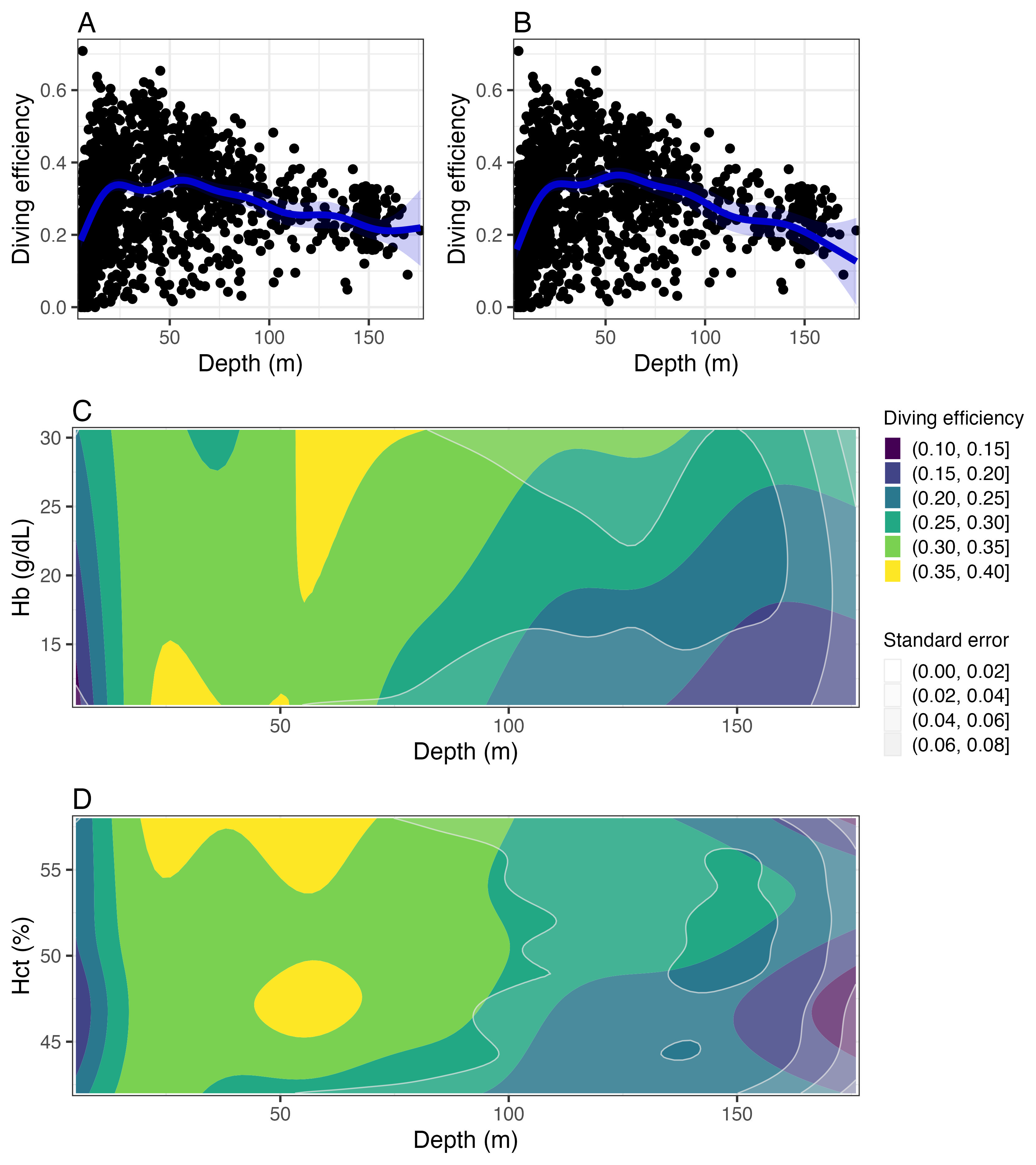

**Figure S4.** Slices of predictions from the two GAMMs that include only one oxygen index. Panels A and B display how the predicted dive efficiency changes with the maximum depth of a dive, for the model with only Hb and Hct respectively. The points show the observed values. Panel C shows how dive efficiency is predicted to change with both depth and Hb for the model that only has Hb as an oxygen index, while panel D shows predicted change with both depth and Hct for the model that only has Hct as an oxygen index. For both panels, the color represents the dive efficiency value and the grey bands represent the standard error. The areas with larger standard error values (i.e., more opaquely shaded) should be interpreted more cautiously. A total of 19 penguins were included in this analysis (5 from Race Point and 14 from Pebble Island).

**Supplement 4. Generalized additive mixed model with a beta distribution**

The residuals of the final GAMM relating diving efficiency with maximum depth, the three indices of oxygen storage and carrying capacity, and colony displayed some evidence of heteroskedasticity (Fig. S5), which can be mostly attributed to the fact that we are modelling proportion data (diving efficiency by definition can only range between 0 and 1) with a normal error distribution. While one should model proportion data with a more appropriate distribution, there are limitations to what can be easily implemented in R. Here, we show that the main results remain unchanged if we use a beta distribution, which is more appropriate for proportion data. We note, however, that using the beta distribution does not allow for diving efficiency values to be exactly zero, and thus we had to add 1e-14 to all zero values. In addition, as far as we are aware, we cannot apply the shrinkage method to select among covariates in R, thus we fit the full model without shrinkage.

The results from the GAMM with the beta distribution are very similar to the results of the final GAMM presented in the main text. In both GAMMs, the only terms that are significant are colony, depth, and the interactions between depth and Hct and depth and Hb (Tables 2 and S3). In both GAMMs, diving efficiency was highest at intermediate depths (Figs. 1A and S6A), and diving efficiency increased with Hb mainly for deeper dives (Figs. 1B and S6B). The complex relationship between diving efficiency and the interaction between Hct and maximum depth also had almost identical patterns (Figs. 1C and S6C). The residuals of the GAMM with a beta regression no longer displayed evidence of heteroskedasticity (Fig. S7).

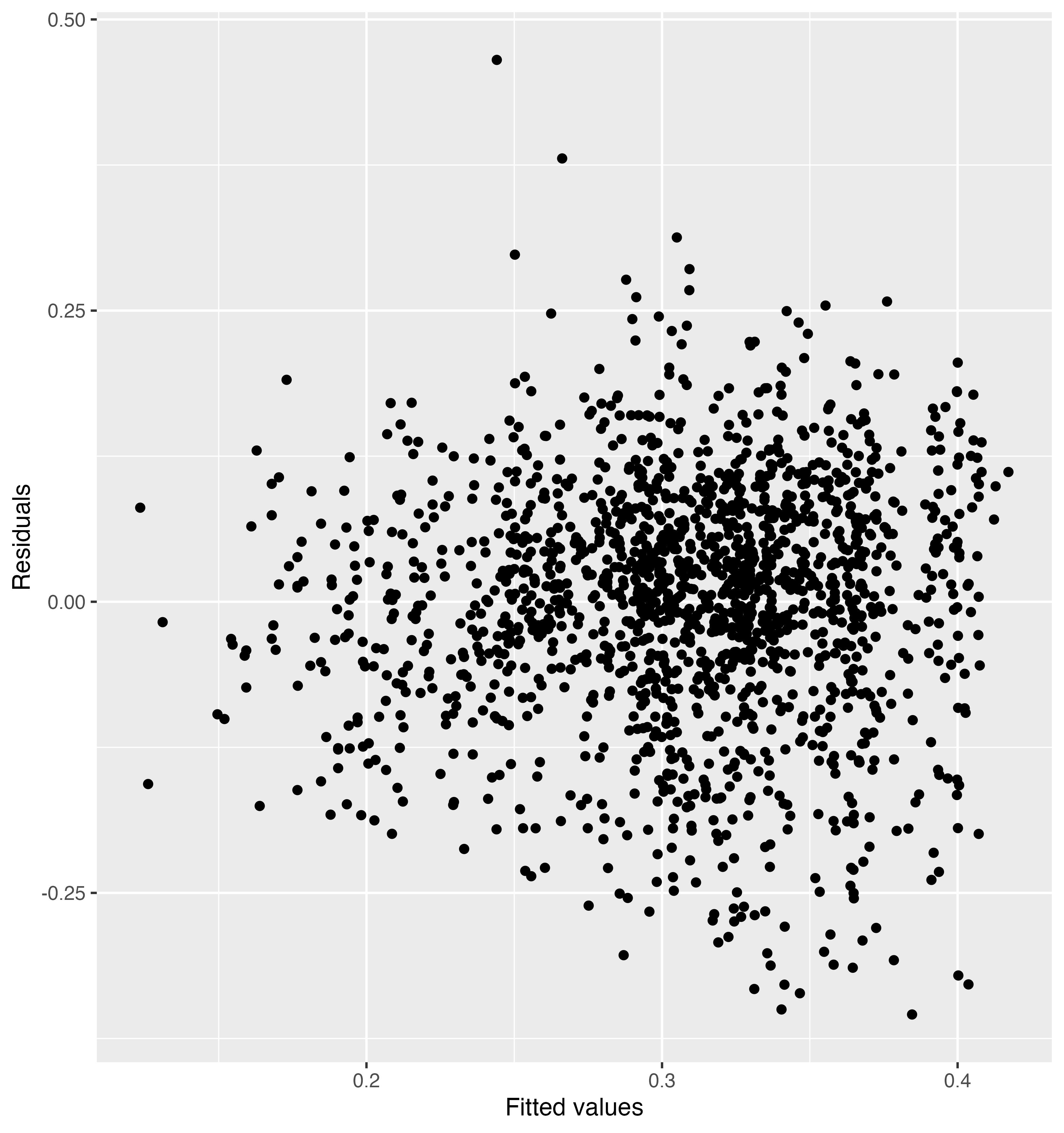

**Figure S5.** Residuals of the final GAMM presented in the main text (i.e., GAMM with normal distribution) against the fitted values. Notice the funnel shape pattern, which shows that variance increases with increasing fitted values.

**Table S3.** Parameter estimates and test statistic values for the GAMM for dive efficiency that uses a beta distribution rather than a normal distribution. A total of 19 penguins were included in this analysis.

| Term | Parametric coefficient | | | | Significance of smooth terms | | |
| --- | --- | --- | --- | --- | --- | --- | --- |
|  | Estimate | SE | *t* | p-value | EDF | F | p-value |
| Intercept | -0.887 | 0.029 | -30.917 | < 0.001 |  |  |  |
| Colony (Race point) | 0.152 | 0.064 | 2.391 | 0.017 |  |  |  |
| s(Depth) |  |  |  |  | 7.974 | 32.608 | < 0.001 |
| s(Mass) |  |  |  |  | 1.000 | 0.026 | 0.873 |
| s(Hct) |  |  |  |  | 1.000 | 2.528 | 0.112 |
| s(Hb) |  |  |  |  | 1.000 | 0.211 | 0.646 |
| ti(Depth, Mass) |  |  |  |  | 1.000 | 3.118 | 0.078 |
| ti(Depth, Hct) |  |  |  |  | 10.685 | 3.892 | < 0.001 |
| ti(Depth, Hb) |  |  |  |  | 9.721 | 4.979 | < 0.001 |

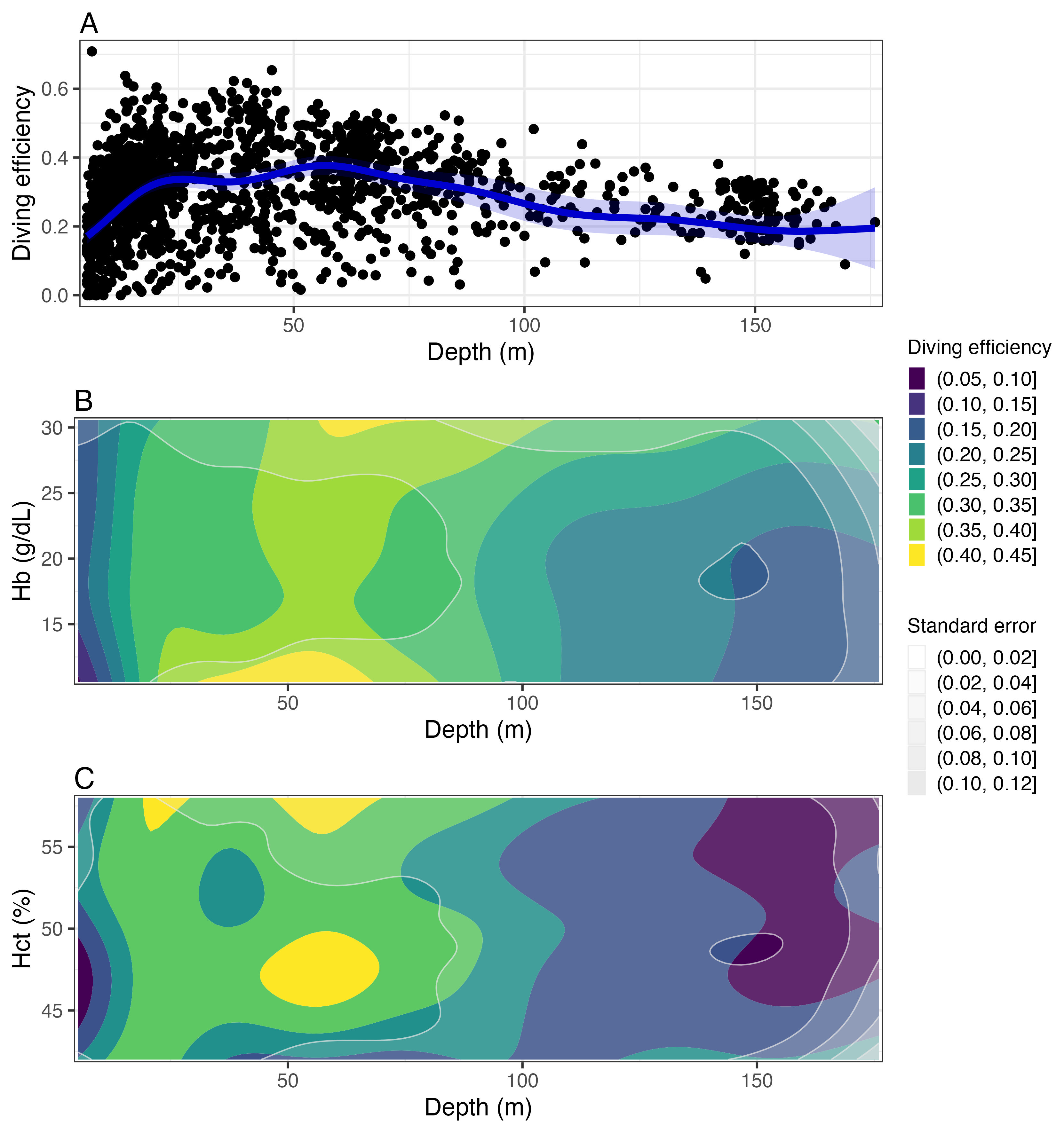

**Figure S6.** Slices of predictions from the GAMM with the beta distribution. Panel A displays how the predicted diving efficiency changes in response to varying maximum depth values of a dive. The blue band represents the 95% confidence intervals. The points show the observed values. Panel B shows how diving efficiency is predicted to change with both depth and Hb, while panel C shows how diving efficiency is predicted to change with both depth and Hct. For both these panels, the colour represents the diving efficiency value (yellow representing highest efficiency) and the grey bands represent the standard error (more opaque grey representing higher standard error). The areas with larger standard error values (i.e., more opaquely shaded) should be interpreted more cautiously. A total of 19 penguins were included in this analysis (5 from Race Point and 14 from Pebble Island).

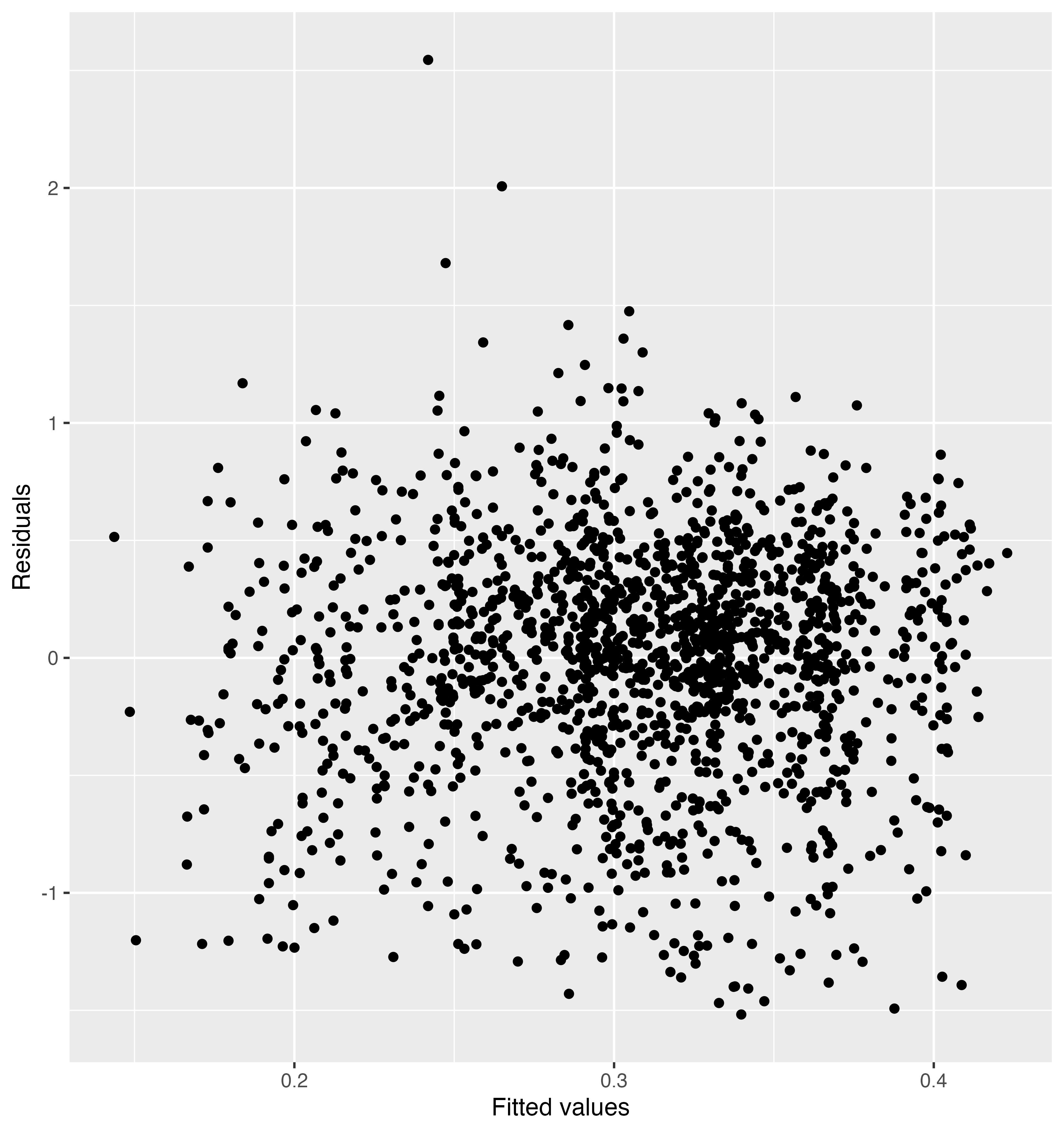

**Figure S7.** Residuals of the GAMM with the beta distribution against the fitted values. Notice that there are no funnel shape patterns and thus that the variance appears to be homogenous across fitted values.

**Supplement 5. General characteristics of the diving, foraging effort, and oxygen storage and carrying capacity indices**

Over the 40 days prior to capture, the TDRs recorded 124,682 dives (> 5 m depth) and 113,495 potential foraging dives with a post-dive surface interval < 200 s. The tagged penguins dove over a wide range of depths, but there were peaks in diving activity centred around a few depths: <10 m, 40 m, 60 m, and 150 m (Fig. S8). Penguins spent an average of 130.0 ± 60.8 s submerged, with the maximum dive time recorded of 520 s occurring at a depth of 174.5 m, and an average of 56.6 ± 32.1 s in the bottom phase of dives, with the maximum bottom time of 228 s occurring at 45.3 m. These penguins had ranging oxygen storage and carrying capacity indices: mass: 6.8 ± 0.6 kg, 5.8 – 7.9 kg; Hb: 20.3 ± 4.9 g/dL, 10.6 – 30.6 g/dL; Hct: 49.1 ± 4.5 %, 42 – 58 %.

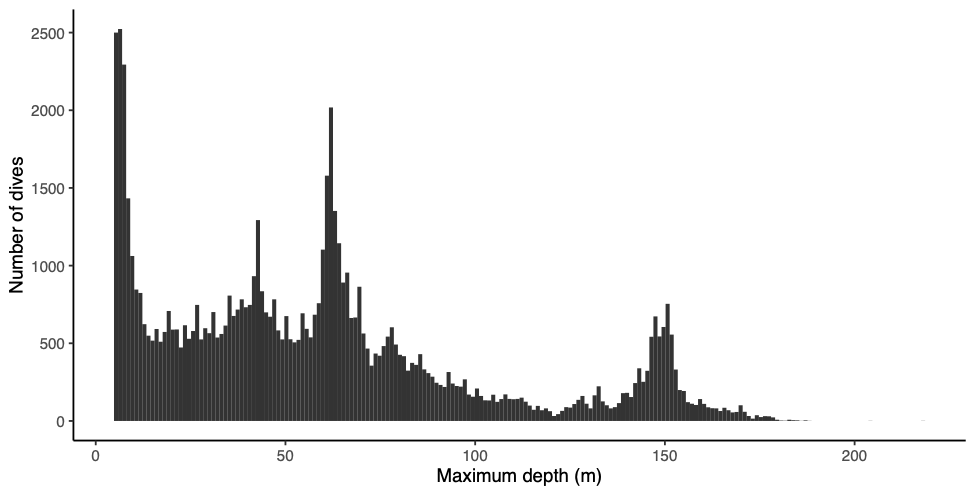

**Figure S8.** Histogram of the maximum depths of the dives performed by 19 penguins over 40 days (5 Race Point, 14 Pebble Island, Falkland Islands).

As just five Race Point penguins were recaptured with successful blood samples and none of the penguins at this colony laid an egg before the departure date of the observers, we decided to assess only the Pebble Island colony (n=14) for connections between foraging effort, oxygen stores, and reproductive status. Behavioural observations resulted in classification of 4 individuals as non-breeders (NB), 6 breeders (B), and 4 early laying (L) individuals, with 8 males (1 NB, 4 B, and 3 L), 5 females (3 NB, 1 B, and 1 L), and one penguin of undetermined sex (1 B). Over the 40 days pre-breeding, these penguins spent an average of 23.2 ± 4.3 days at sea (14.4 – 30.2 days). Penguins went on 13 ± 4 trips, for which mean trip duration lasted 2.6 ± 3.5 days at sea. Many individuals performed long trip durations (e.g., 43 % of individuals went on trips longer than 10 days). The long trips tended to be at the beginning of the 40 days period before recapture. While 69.8 % of all trips lasted under one day, 9.0 % lasted over five days in length, with the longest trip spanning 24.3 days. Long trip durations such as these have been rarely reported for gentoo penguins and may be evidence of a pre-breeding exodus (e.g., Mallory et al. 2008).

**Supplement 6. Results from the best models relating foraging measures to covariates**

**Table S4.** Parameter estimates and test statistic values for the best models relating vertical distance travelled and time at sea with covariates. The best model for average trip duration was the null model, and is thus not presented.

|  | Term | Estimate | SE | *t* | p-value |
| --- | --- | --- | --- | --- | --- |
| *Vertical distance travelled* | | | | | |
|  | Intercept | 149.526 | 191.346 | 0.781 | 0.451 |
|  | Hb | 19.955 | 8.627 | 2.313 | 0.041 |
| *Time at sea* | | | | | |
|  | Intercept | 25.220 | 1.058 | 23.834 | <0.001 |
|  | Breeding status (Breeding) | -0.978 | 1.420 | -0.689 | 0.507 |
|  | Breeding status (Early laying) | -5.518 | 1.496 | -3.687 | 0.004 |

**Supplement 7. Comparison in breeding status between tagged and untagged penguins**

While we did not find any significant difference in the average values of Hb and Hct between tagged and untagged penguins (Hb: t_62.863_ = -0.524, p-value = 0.774, n = 66; Hct: t_63.633_ = 1.480, p-value = 0.144, n = 66; Fig. S9A-B), the tagged penguins were on average 5% lighter than the untagged penguins (tagged penguins: 6.9 kg ± 0.6, untagged penguins: 7.3 kg ± 0.5; t_66.498_ = 2.812, p-value = 0.006, n = 70; Fig. S9C). The additional drag created by the tags can affect the foraging success of penguins (Wilson et al. 2015), and other diving seabirds have shown declines in body mass associated with tagging (Evans et al. 2020). While the differences were not statistically significant (Pebble Island: χ^2^ = 1.167, simulated p-value = 0.604, n = 39; All: χ^2^ = 1.26, simulated p-value = 0.548, n = 70), we note that compared to the untagged penguins, there was a higher percentage of tagged penguins that did not participate in breeding and a lower percentage of tagged penguins that laid early (Table S5). Tagging has been shown to affect the breeding success of penguins (e.g., Beaulieu et al. 2010), and our results may suggest that our tagging may have had a negligible effect that is not large enough to be detected with our sample size. Taken together, our results suggest that it is unlikely the relationships we found with Hb and Hct are affected by tagging, but that tagging could have reduced the mass of the penguins and may affect their behaviour. Such impacts should be further considered before conducting other tagging studies with gentoo penguins.

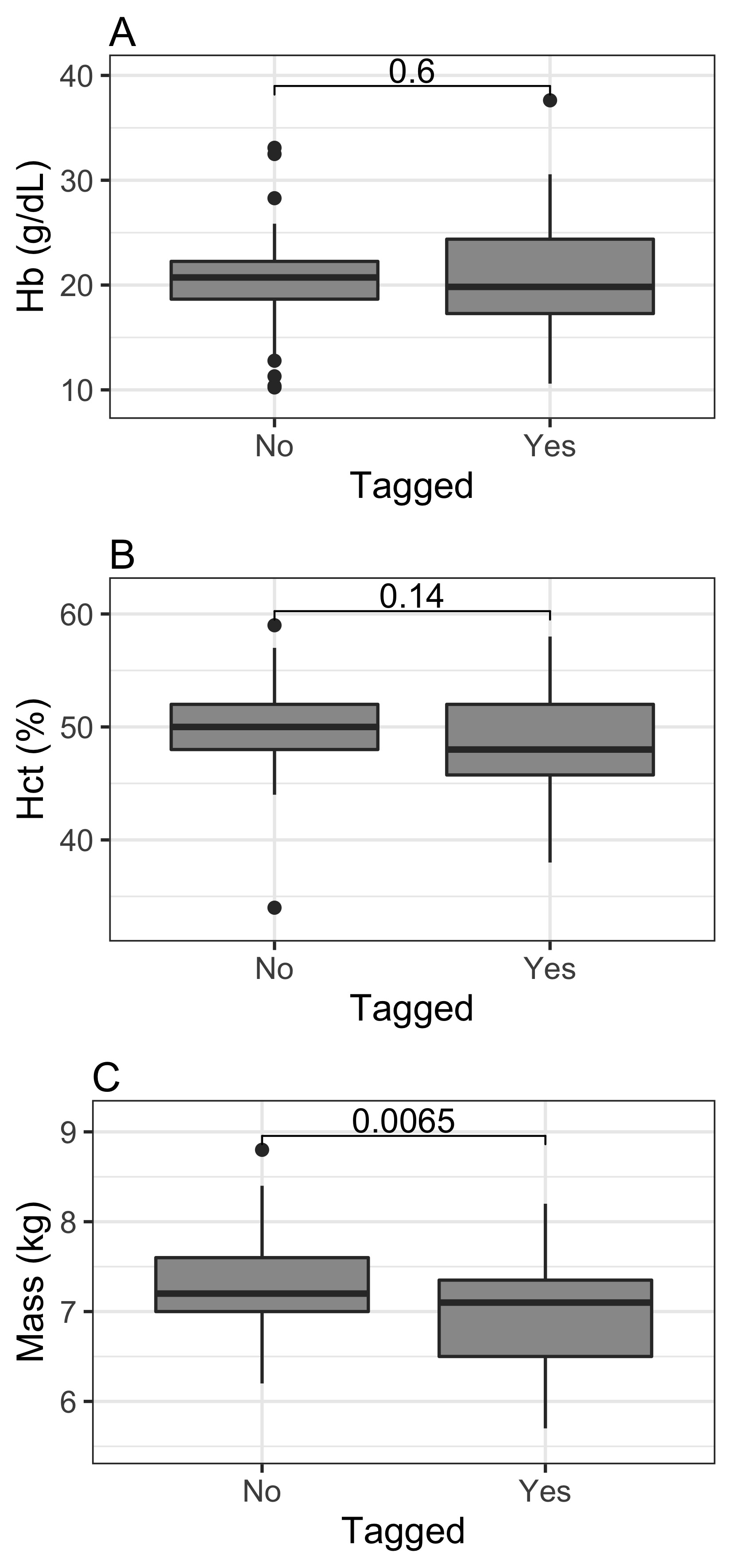

**Figure S9**. Differences between tagged and untagged penguins in terms of Hb (panel A), Hct (panel B), and mass (panel C). The values presented in the panels are the p-values associated with the Welch two-sample t-test comparing the tagged and untagged penguins. We have Hb and Hct data for 32 tagged and 34 untagged penguins (panels A-B) and mass for 35 tagged and 35 untagged penguins (panel C).

**Table S5.** Contingency table presenting the number of tagged and untagged penguins in each of the breeding categories: non-breeding, breeding, and early-laying. The percentages represent how many tagged or untagged penguin are in each of the breeding categories, and is presented to facilitate the comparison between tagged and untagged penguins.

|  | Race Point | | Pebble Island | | Total | |
| --- | --- | --- | --- | --- | --- | --- |
|  | Tagged | Untagged | Tagged | Untagged | Tagged | Untagged |
| Non-breeding | 12 (75%) | 10 (67%) | 5 (26%) | 3 (15%) | 17 (49%) | 13 (40%) |
| Breeding | 4 (25%) | 5 (33%) | 9 (47%) | 9 (45%) | 13 (37%) | 14 (37%) |
| Early laying | 0 (0%) | 0 (0%) | 5 (26%) | 8 (40%) | 5 (14%) | 8 (23%) |

**References**

Evans TJ, Young RC, Watson H, Olsson O, Åkesson S (2020). Effects of back-mounted biologgers on condition, diving and flight performance in a breeding seabird. Journal of Avian Biology 51:e02509.

Handley JM, Thiebault A, Stanworth A, Schutt D, Pistorius P (2018) Behaviourally mediated predation avoidance in penguin prey: in situ evidence from animal-borne camera loggers. Royal Society Open Science 5:171449.

Houstin A, Zitterbart DP, Winterl A, Richter S, Planas-Bielsa V, Chevallier D, Ancel A, Fournier J, Fabry B, le Bohec C (2022) Biologging of emperor penguins—Attachment techniques and associated deployment performance. PLoS ONE 17:e0265849.

Mallory ML, Forbes MR, Ankney CD, Alisauskas RT (2008) Nutrient dynamics and constraints on the pre-laying exodus of high Arctic northern fulmars. Aquatic Biology 4:211–223.

Marra G, Wood SN (2011) Practical variable selection for generalized additive models. Computational Statistics and Data Analysis 55:2372–2387.

Wilson RP, Pütz K, Peters G, Culik B, Scolaro JA, Charrassin J-B, Ropert-Coudert Y (1997) Long-term attachment of transmitting and recording devices to penguins and other seabirds. Wildlife Society Bulletin 25:101–106.

Wilson RP, Sala JE, Gómez-Laich A, Ciancio J, Quintana F (2015) Pushed to the limit: food abundance determines tag-induced harm in penguins. Animal Welfare 24:37–44.

Wood SN (2011) Fast stable restricted maximum likelihood and marginal likelihood estimation of semiparametric generalized linear models. Journal of the Royal Statistical Society (B) 73:3-36.
